# Hydrothermal origin of metabolic phosphorylation

**DOI:** 10.64898/2025.12.19.695421

**Authors:** Manon L. Schlikker, Nadja K. Hoffmann, Sabine Metzger, Jorge Moral-Pombo, Harun Tüysüz, William F. Martin

## Abstract

Phosphate is central to modern bioenergetics and to all theories for the origin of life. How phosphate entered metabolism is unknown, though microbial physiology and geochemical environments can provide important clues. Some bacteria obtain electrons and energy from phosphite (HPO_3_^2–^), a reduced form of phosphate (HPO_4_^2–^), that naturally occurs in serpentinizing (H_2_-producing) hydrothermal systems. Here we show that the insoluble, solid-state catalyst native palladium, which is naturally deposited in serpentinizing hydrothermal systems, catalyzes the oxidation of phosphite to phosphate and H_2_ in water at 25–100 °C in a highly exergonic reaction. Palladium awaruite (Pd_x_Ni_3_Fe), a common form of Pd^0^ in serpentinizing vents, also catalyzes phosphite-dependent phosphorylation. Phosphite oxidation over Pd^0^ generates a reactive but so far unidentified chemical intermediate, possibly metaphosphate, [PO_3_]^−^, that readily phosphorylates hydroxyl moieties in glycerol, ribose, glucose, serine and cytidine at 25–100 °C in 2–72 h. The same conditions also generate (i) phosphoanhydride bonds in pyrophosphate, polyphosphates and ADP, (ii) the phosphoramidate bond in phosphocreatine, (iii) and the acyl phosphate bond in acetyl phosphate, which is obtained overnight at 25 °C with 8% yield. The reactions proceed without sulfur, excluding thioester or metal sulfide intermediates. Phosphite-dependent phosphorylations under serpentinizing hydrothermal vent conditions are facile. They identify a natural, geochemical source of prebiotic phosphorylation and a novel source of metabolic energy at origins. The central role of phosphate in bioenergetics, metabolism, and nucleic acids could reflect metal-catalyzed, redox chemistry of phosphorus in the environment where metabolism (and life) arose.

## Introduction

Theories for the origin of life hinge upon mechanisms for the incorporation of phosphate into organic compounds [1–3]. In genetics first theories [4, 5], phosphate is an essential structural component of nucleic acid synthesis and replication, as RNA is ∼30% phosphate by weight. In metabolism first theories [6–9], phosphate is a transient carrier of metabolic energy. It serves as a universal energy currency of cells, reacting with organic or organophosphate substrates to form metastable bonds [1] that release free energy upon hydrolysis. Phosphorylation activates molecules for further synthesis and couples energy release to otherwise endergonic reactions, pushing chemical reactions in the life process forward [10, 11].

Possible environmental sources and states of phosphorus at origins are still debated [12, 13]. In the biosynthesis of amino acids, nucleotides, and cofactors of modern cells, phosphorus exists exclusively as phosphate [14, 15]. Phosphonates, compounds with a C–P bond, are synthesized and utilized by microbes [16, 17], but not for ATP or nucleic acid synthesis [16, 18]; they are synthesized from phospho*enol*pyruvate (a phosphate ester) without redox reactions [19] and they release phosphate upon degradation [18]. The most common phosphorus mineral in nature, calcium phosphate (apatite), is poorly soluble [20, 21]. Phosphate is not readily incorporated into organic compounds via prebiotic syntheses without enzymes, requiring the use of non-physiological conditions including oven-heating to dryness in the presence of condensing agents such as ammonia and urea [22, 23], incubation with phosphate in pure formamide [24], strong non-physiological electrophiles such as cyanoacetylene, cyanogen, cyanate, cyanoformamide, cyanamide [25], or highly reactive forms of phosphorus [26] such as P_4_O_10_ [13] These chemical impasses to phosphate incorporation are also known as the “phosphate problem” [2], which has a long history [27] and persists to the present, as Weller *et al.* [13] recently summarized: “*a fundamental, universally accepted mechanism how early Earth could have provided phosphate for prebiotic chemistry remains unknown*.” Yet phosphorus was absolutely essential for metabolism, information, heredity and energy at origins, indicating that facile routes of prebiotic phosphorylation must have existed on the early Earth. How to find them?

We reasoned that microbial physiology [28] and the chemically reactive environments of hydrothermal vents [29] in particular serpentinizing (alkaline, H_2_ producing) hydrothermal vents [30] can point to possible natural sources of phosphorylation. Bernhard Schink and colleagues isolated bacteria that can use phosphite as their sole phosphorus source, converting it to phosphate via enzymatic phosphite oxidation [31–33], prompting the suggestion by Wolfgang Buckel [34] that phosphite could have generated acyl phosphates via phosphite oxidation at a biochemical origin. Though phosphite is vastly more soluble than phosphate, reports for its occurrence in modern natural environments are rare [17, 35]. Yet phosphite-oxidizing genes are extremely common among modern microbes, with up to 1.5% of sequenced prokaryotes being able to utilize phosphite [36]. Phosphite was furthermore present in Archaean oceans [37] and in Archaean rocks [35].

Microbial genomes obtained from serpentinizing hydrothermal vents have an increased frequency of phosphite-oxidizing genes [38–40]. Moreover, Pasek *et al.* [41] measured the frequency of different phosphorus species in rock samples from formations altered by serpentinization and found that between 20% to 50% of the total phosphorus in some samples was phosphite, the remainder being phosphate. Why does phosphite occur in serpentinized rocks? Serpentinization is a geochemical process [42–44] in which water circulating 1–5 km in the Earth’s crust is reduced by reactions with Fe(II) minerals, generating very high H_2_ concentrations (often >10 mM) and very alkaline (pH 9–12) effluent [39]. The result is that the fluids in serpentinizing systems generate potentials up to –900 mV [45], sufficient to reduce dissolved transition metals to the native (zero valent, metallic) state [46] and, by inference, reduce phosphate to phosphite [41], as the midpoint potential of the HPO_4_^2-^ to HPO_3_^2-^ redox couple is –690 mV [31]. While phosphite itself has not yet been reported in the effluent of hydrothermal vents, its presence in serpentinized rocks [41] and the frequency of phosphite oxidizing genes in microbes of serpentinizing systems [38–40] prompted us to investigate its ability to phosphorylate organic compounds using conditions (aqueous, alkaline) and catalysts germane to serpentinizing vents.

As catalysts for phosphite activation, we focused on native transition metals, because (i) native metals are naturally deposited in serpentinizing hydrothermal systems [46] and because (ii) under aqueous conditions, native transition metals have been found to readily convert H_2_ and CO_2_ to organic acids [47–49], convert NH_3_ and organic acids to amino acids [50, 51], convert H_2_ and CO_2_ to long chain hydrocarbons [52], to functionally interact with NAD^+^ [53, 54], pyridoxal [51], and ferredoxin [55], and to reduce double bonds of biochemical compounds [56]. In an initial screen [15] we found that Pd^0^, a metal that occurs in the native state in serpentinizing hydrothermal vents [57–59], was more effective than Fe^0^, Co^0^, Ni^0^ or Ni_3_Fe in the oxidation of HPO_3_^2–^ to HPO_4_^2–^. Hence, we probed the ability of HPO_3_^2-^ over Pd^0^ in water to phosphorylate biochemicals under prebiotic conditions.

## Results

Mao *et al.* [33] characterized the enzyme that the bacterium *Phosphitispora fastidiosa* uses to access electrons and generate ATP from phosphite: AMP-dependent phosphite dehydrogenase (AdpA). AdpA catalyzes the reaction AMP + HPO_3_^2–^ + NAD^+^ ® ADP + NADH, whereby ADP is subsequently converted to ATP via adenylate kinase: 2ADP → ATP + AMP. The enzyme does not harbor a metal cofactor. The oxidation of HPO_3_^2–^ via the reaction HPO_3_^2–^ + H_2_O → HPO_4_^2–^ + H_2_ is exergonic with Δ*G*_0_′ = –46.3 kJ·mol^−1^ [60]. In water, Pd^0^ oxidizes HPO_3_^2–^ to HPO_4_^2–^, generating H_2_ during the reaction. In the presence of AMP, the palladium catalyzed reaction is AMP + HPO_3_^2–^ ® ADP + H_2_, with 4.3% conversion after 2 h and 5.9% conversion after 18 h at 50 °C (**Fig. 1A**). The ADP product was identified using ^1^H NMR (black line, **Fig. 1B**) by comparison to the library standard (blue area, **Fig. 1B**) and using ^31^P NMR, with specific peaks for adenosine 5’-diphosphate at –6 and –10.4 ppm (**Fig. 1C**). ADP yields decrease with temperature, with breakdown to AMP + P_i_ and adenosine (not shown). Though far less efficient than AdpA, Pd^0^ (as Pd^0^ on carbon, Pd/C) replaces the AdpA enzyme for phosphite to phosphate (in ADP) oxidation and uses H^+^ instead of NAD^+^ as the electron acceptor, generating H_2_ instead of NADH.

**Figure 1.**
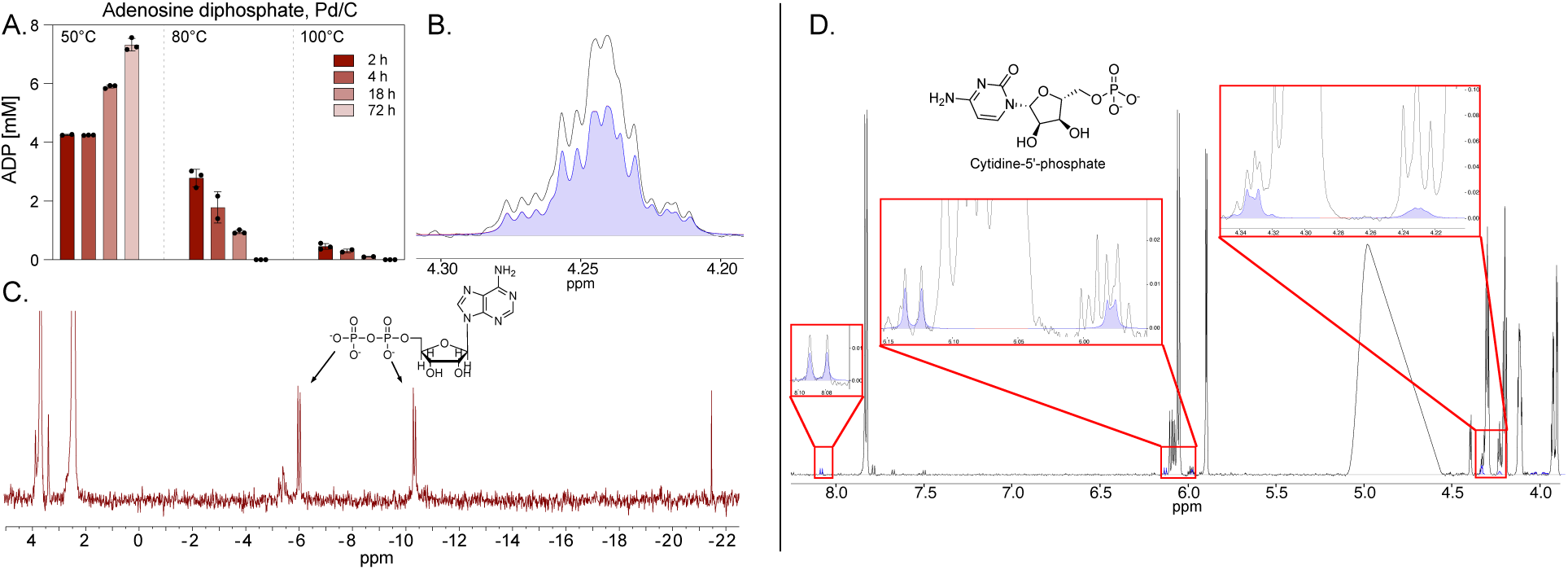
Palladium-catalyzed phosphorylation of AMP and cytidine using phosphite. **(A)** Formation of adenosine diphosphate (ADP) from AMP (100 mM) and phosphite (200 mM) in the presence of Pd/C (1.5 mmol) under 5 bar Ar at pH 9, reaction volume 1.5 ml. Reaction temperatures were 50 °C, 80 °C, and 100 °C stopped after 2 h, 4 h, 18 h, and 72 h. ADP concentrations were estimated by ^1^H NMR integration (**Fig. S1, Table S1**). **(B)** ^1^H NMR spectrum (black) of the 50 °C, 72 h reaction, showing a diagnostic ADP signal at ∼4.25 ppm; purple overlay represents the internal library reference. **(C)** ^31^P NMR spectrum of the same sample, showing ADP peaks at -6 ppm and -10.4 ppm. **(D)** ^1^H NMR spectrum of cytidine (100 mM) reacted with phosphite (200 mM) under identical conditions (Pd/C, 50 °C, 72 h, pH 9, 5 bar Ar), showing signals consistent with the formation of cytidine 5′-monophosphate (CMP; purple insets). The presence of cytidine phosphate was further supported by ESI–LC–MS analysis, and the corresponding raw data are provided in **Fig. S2** and **Table S2**.

In metabolism, the 5ʹ-phosphate of nucleotides stems from 5-phosphoribosyl-1-pyrophosphate, while prebiotic-type nucleotide syntheses typically involve nucleoside phosphorylation. Phosphite over Pd^0^ phosphorylates cytidine at the 5ʹ position to cytidine 5ʹ-monophosphate at 30 °C in 72 h, albeit at low yields of ca. 0.37% in ^1^H NMR spectra (**Fig. 1D**). Though our yields are an order of magnitude lower than those obtained using phosphate to phosphorylate uridine in the presence of molar concentrations of cyanogen, cyanoformamide, cyanate, cyanamide, thioformate, ethylisocyanide, and a carbodiimide [22, 25], our reaction requires no organic additives, only phosphite and water, plus an effective solid state transition metal catalyst. Because readers unfamiliar with heterogeneous catalysis might be concerned about the concentration of Pd^0^ in our reactions, we clarify that our Pd^0^ catalyst is not dissolved; it is in the solid phase and has a concentration of zero, the same concentration as glass from the walls of vials in which our reactions are performed. AMP phosphorylation using 75 mM HPO_3_^2–^ over Pd/C, instead of 200 mM, 50 °C, 72 h, generated a yield of 0.7% (**Fig. S1)**.

### Hydroxyl groups

In modern metabolism, sugar phosphates are essential for the biosynthesis of amino acids, nucleic acids, reserve polysaccharides and cell wall components. In autotrophs that use the acetyl-CoA pathway, ribose 5-phosphate is generated through gluconeogenesis [61] and the oxidative pentose phosphate pathway [62]. Non-enzymatic phosphorylation of sugars can be attained through heating with phosphate and condensing agents such as cyanogen or cyanamide [25], but the existence of highly reactive nitrile-containing compounds such as cyanogen or cyanamide on the early Earth is uncertain at best [63] while the existence of serpentinizing systems on the early Earth is certain [64–66]. Pd/C catalyzes the phosphorylation of ribose to ribose 5-phosphate at 0.6% yield (**Fig. 2A**). The yield is comparable to that obtained through heating ribose and phosphate with cyanamide [25, 67], a procedure that is also used to phosphorylate glucose [25, 68]. Phosphite with Pd/C generates glucose 6-phosphate (**Fig. 2B**) at yields of 1.1% (at least 10-fold lower than the classical cyanogen method), but in pure water, without the requirement for addition of cyanogen gas or cyanamide [67, 68], and at a concentration of 1.1 mM, slightly lower than the physiological concentration of glucose 6-phosphate in *E. coli* grown on glucose, 1.5 mM [69].

**Figure 2.**
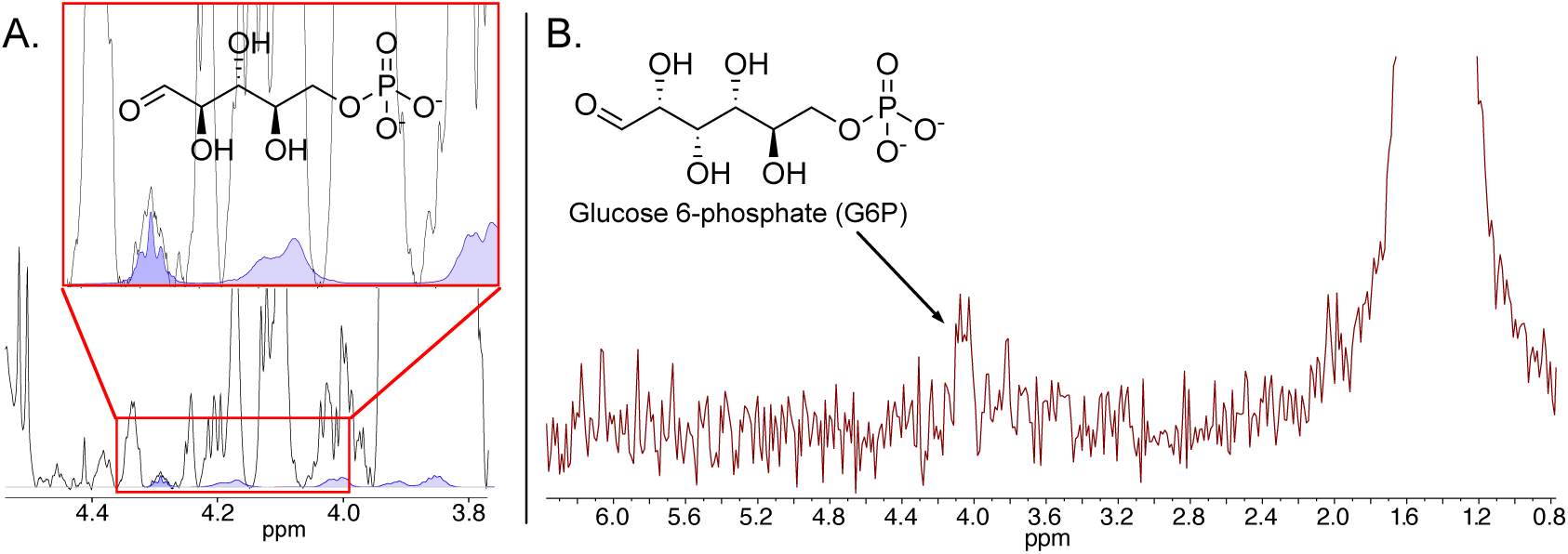
Pd/C-catalyzed phosphorylation of ribose and glucose using phosphite as phosphate donor. **(A)** ^1^H NMR spectrum of the 50 °C, 72 h ribose (100 mM) and phosphite (200 mM) reaction in the presence of Pd/C (1.5 mmol) under 5 bar Ar at pH 9 in a total reaction volume = 1.5 mL. R5P was detected only at 50 °C after 18 h and 72 h. Showing a characteristic ribose 5-phosphate (R5P) signal at ∼4.28 ppm (assignment supported by ESI–LC–MS analysis; corresponding raw data in **Fig. S3, Table S2**). **(B)** ^31^P NMR spectrum of the glucose phosphorylation reaction (100 mM glucose, 200 mM phosphite, 1.5 mmol Pd/C, 40 °C, 18 h) under Ar at pH 9, with a phosphate resonance consistent with G6P at ∼3.9 ppm (**Fig. S4**).

Although it is possible, if not probable, that the lipids of bacteria and archaea were synthesized by enzymatic processes in early evolution [9, 70], glycerol phosphate was required for phospholipid synthesis. Glycerol phosphorylation has been reported using schreibersite, Fe_3_P, a highly reactive phosphide that occurs in meteorites [71]. With HPO_3_^2–^ and Pd/C in water, glycerol is readily phosphorylated with 0.8% yield after 4 hours and 1.7% yield after 72 hours at 50 °C, with slightly lower yields at higher temperatures (**Fig. 3A**). The hydroxyl group of serine is very efficiently phosphorylated over Pd/C with ca. 49.5% yield after 18 h and 50 °C (**Fig. 3D**), and lower yields at higher temperatures. Using Pd^0^ nanoparticles, without carbon support, serine phosphorylation yield is generally lower, but increases with time and temperature (**Fig. 3E**).

**Figure 3.**
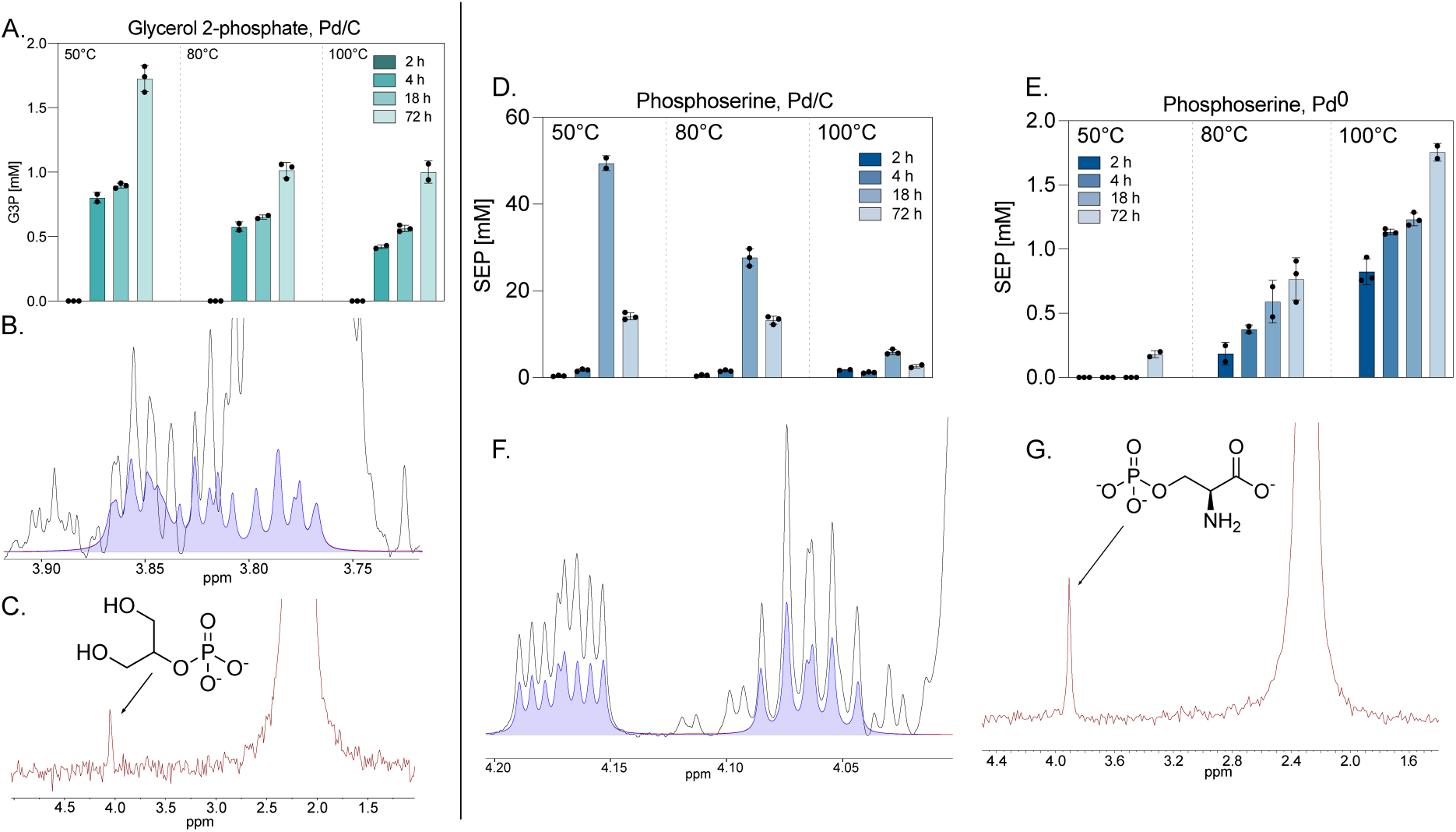
Phosphorylation of glycerol and serine by phosphite in water. (A–C) Glycerol 2-phosphate (G2P) formation from 100 mM glycerol and 200 mM phosphite over 1.5 mmol Pd/C in water under Ar at pH 9, reaction volume 1.5 ml. **(A)** G2P yield at 50 °C, 80 °C, and 100 °C after 18 h and 72 h (**Fig. S5, Table S1**). **(B)** ^1^H NMR spectrum of the 50 °C, 72 h reaction (black), overlaid with internal library standard (purple). **(C)** ^31^P NMR spectrum of the same sample shows a characteristic peak for G2P. **(D–G)** Phosphorylation of 100 mM serine under identical conditions. **(D)** Serine phosphorylation yield using Pd/C (**Fig. S6, Table S1**). **(E)** Yield with native Pd^0^ nanoparticles (**Fig. S7, Table S1**). **(F)** ^1^H NMR spectrum of Pd/C reaction after 18 h at 50 °C (black) and internal standard (purple). **(G)** ^31^P NMR spectrum of the same sample confirms the presence of phosphoserine.

### High-energy bonds

A key to phosphate in energy metabolism is its ability to form generally long, readily attacked (and hydrolyzed) metastable bonds with various moieties [72]. Hydrolysis of such bonds releases free energy that can be harnessed for phosphorylation of other compounds or enzymatically coupled to endergonic reactions, rendering the coupled reaction exergonic [11]. The free energy of hydrolysis for some bioenergetically relevant reactions under standard physiological conditions (25 °C, pH 7) is given in **Table 1**. In cells, high-energy phosphate bonds are synthesized as ATP through the stoichiometric coupling of enzymatic reactions to ATP synthesis via substrate-level phosphorylation (SLP) [10, 73] or from chemiosmotic coupling through the harnessing of ion gradients via rotor-stator ATPases [73, 74].

**Table 1.** Free energy of hydrolysis for selected reactions.

| Hydrolysis reaction | $\Delta G_0'$ | Ref. |
| --- | --- | --- |
| PEP + H<sub>2</sub>O → HPO<sub>4</sub><sup>2-</sup> + pyruvate | −62 | 75 |
| 1,3-BPGA + H<sub>2</sub>O → HPO<sub>4</sub><sup>2-</sup> + 3PGA | −52 | 75 |
| HPO<sub>3</sub><sup>2-</sup> + H<sub>2</sub>O → HPO<sub>4</sub><sup>2-</sup> + H<sub>2</sub> | −46 | 60 |
| ATP + H<sub>2</sub>O → AMP + pyrophosphate | −46 | 75 |
| Creatine-P + H<sub>2</sub>O → HPO<sub>4</sub><sup>2-</sup> + creatine | −43 | 75 |
| Acetyl-P + H<sub>2</sub>O → HPO<sub>4</sub><sup>2-</sup> + acetate | −43 | 75 |
| Carbamoyl-P + H<sub>2</sub>O → HPO<sub>4</sub><sup>2-</sup> + carbamate | −39 | 75 |
| Acetyl-CoA + H<sub>2</sub>O → CoASH + acetate | −32 | 76 |
| ATP + H<sub>2</sub>O → HPO<sub>4</sub><sup>2-</sup> + ADP | −31 | 11 |
| Glucose 1-P + H<sub>2</sub>O → HPO<sub>4</sub><sup>2-</sup> + glucose | −21 | 75 |
| Pyrophosphate + H<sub>2</sub>O → 2 HPO<sub>4</sub><sup>2-</sup> | −20 | 75 |
| Glucose 6-P + H<sub>2</sub>O → HPO<sub>4</sub><sup>2-</sup> + glucose | −14 | 75 |
| Glycerol 2-P + H<sub>2</sub>O → HPO<sub>4</sub><sup>2-</sup> + glycerol | −9 | 75 |

Note: The value of $\Delta G_0'$ for the phosphite oxidizing reaction shown has also been reported as ca. $-55 \text{ kJ} \cdot \text{mol}^{-1}$ [77]

Before enzymes, SLP and chemiosmotic harnessing via ATP synthases did not exist. Metabolic reactions are continuous in nature. What was the continuous source of high-energy bonds at metabolic origin? In water, HPO_3_^2–^ over Pd/C phosphorylates acetate, generating acetyl phosphate (**Fig. 4A, B**) with a yield of 8%. The acyl phosphate bond in acetyl phosphate has a free energy of hydrolysis Δ*G*_0_′ = –43 kJ·mol^−1^ (**Table 1**), sufficient to phosphorylate a variety of substrates including ADP to ATP via SLP [11]. In water, HPO_3_^2–^ over Pd/C also phosphorylates creatine, generating the phosphoamidate bond in phosphocreatine (**Fig. 4C**) which also has a free energy of hydrolysis Δ*G*_0_′ = –43 kJ·mol^−1^ (**Table 1**). Phosphocreatine is a phosphagen that is used as a reservoir of high-energy bonds to rapidly synthesize ATP in muscle via the enzyme creatine kinase. A role for creatine and other phosphagens in prebiotic chemical evolution now appears possible. In water, HPO_3_^2–^ over Pd/C also phosphorylates phosphate to form pyrophosphate (**Fig. 4D**). Pyrophosphate is traditionally considered as a primordial energy currency [10], but it has a lower free energy of hydrolysis than glucose 1-phosphate (**Table 1**) and is more likely a currency of irreversibility [78, 79] than energy.

**Figure 4.**
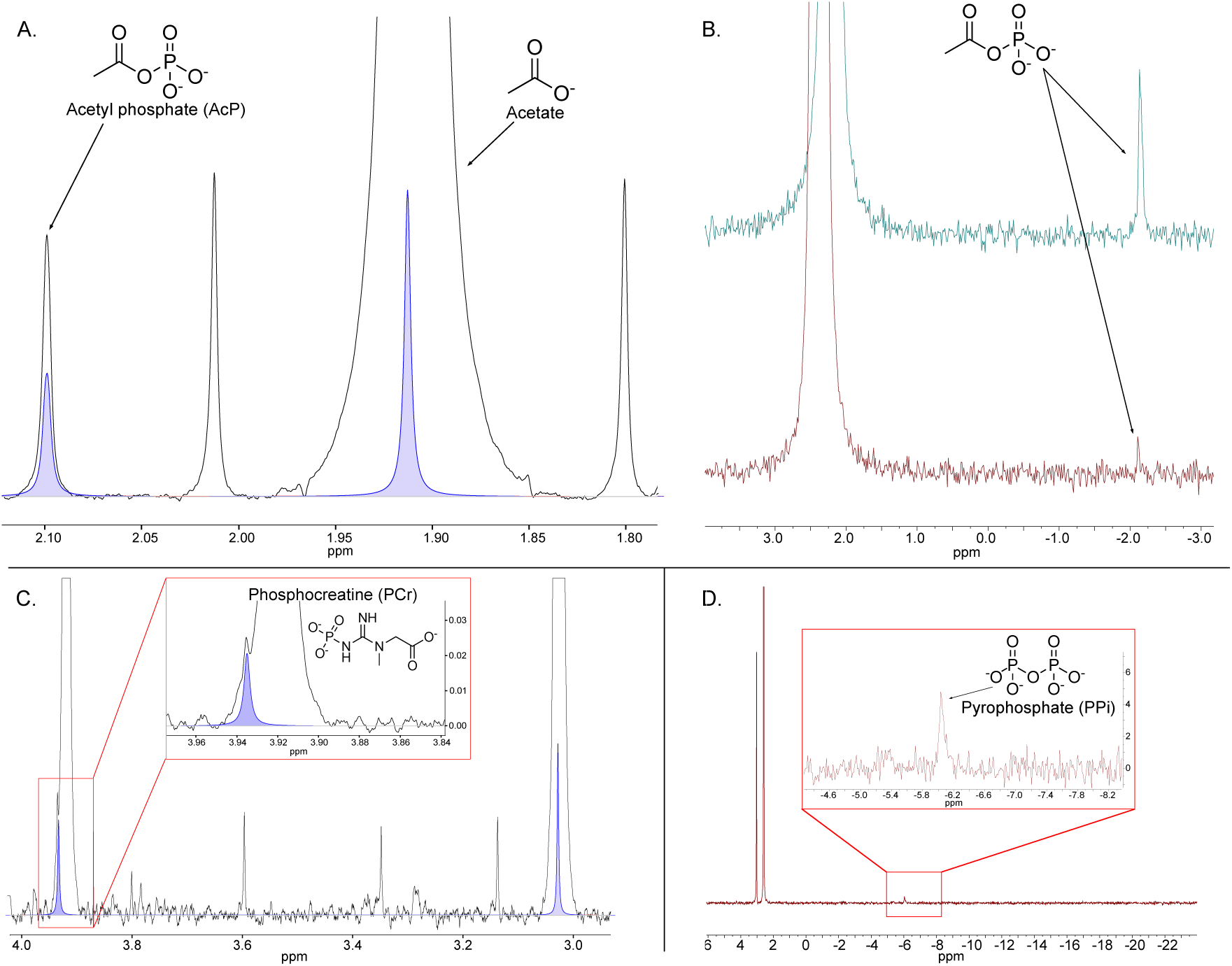
Formation of energy-rich phosphoanhydrides via Pd/C-catalyzed phosphite oxidation. **(A)** ^1^H NMR spectrum of a reaction containing acetate (100 mM) and phosphite (200 mM) with Pd/C (1.5 mmol) in 1.5 ml water under 5 bar Ar at pH 9 after 18 h at 25 °C. Two natural abundance ^13^C acetate satellite peaks appear at 2.02 and 1.8 ppm [80]. **(B)** ^31^P NMR spectra showing AcP identification via co-injection. Top: reaction sample spiked with AcP (cyan); bottom: unspiked reaction mixture (red), confirming identity (**Fig. S8**). **(C)** Formation of phosphocreatine (PCr) from creatine (100 mM) and phosphite (200 mM) under identical conditions with Pd/C (1.5 mmol) at 60 °C for 72 h (^1^H NMR, black trace), confirmed by overlay with Chenomx library reference (purple) (**Fig. S9, Table S2**). **(D)** ^31^P NMR spectrum of a reaction (200 mM phosphite) at 80 °C after 18 h, showing pyrophosphate (PPi) formation. Signals for residual phosphate (Pi) and unreacted phosphite (Pt) are also visible.

The synthesis of pyrophosphate (**Fig. 4D**) suggests that further phosphate phosphorylations are possible with HPO3^2–^ over Pd/C. We observe the synthesis of polyphosphate (**Fig. 5A**) and, surprisingly in our view, trimetaphosphate (**Fig. 5B**). Polyphosphates are very common among bacteria and archaea and are commonly viewed as a potential energy reserves and phosphate storage compounds [81, 82]. That HPO_3_^2–^ can generate polyphosphates over Pd in water suggests that these have been in existence since phosphate was first used as an energy currency. More surprising was the detection of trimetaphosphate, yet the ^31^P NMR signal observed in **Fig. 5B** is identical to that reported by Gull *et al.* [83]. Trimetaphosphate (P_3_O_9_^3–^) is not known from cells but is widely used as a phosphorylating agent in prebiotic chemistry [84]. That trimetaphosphate is formed in addition to a broad range of physiological phosphorylated products suggests that it could have served as a phosphorylating agent in early biochemistry. We cannot presently exclude the possibility that trimetaphosphate might be involved as a transient, undetected intermediate in some of the phosphorylation reactions we report here.

**Figure 5.**
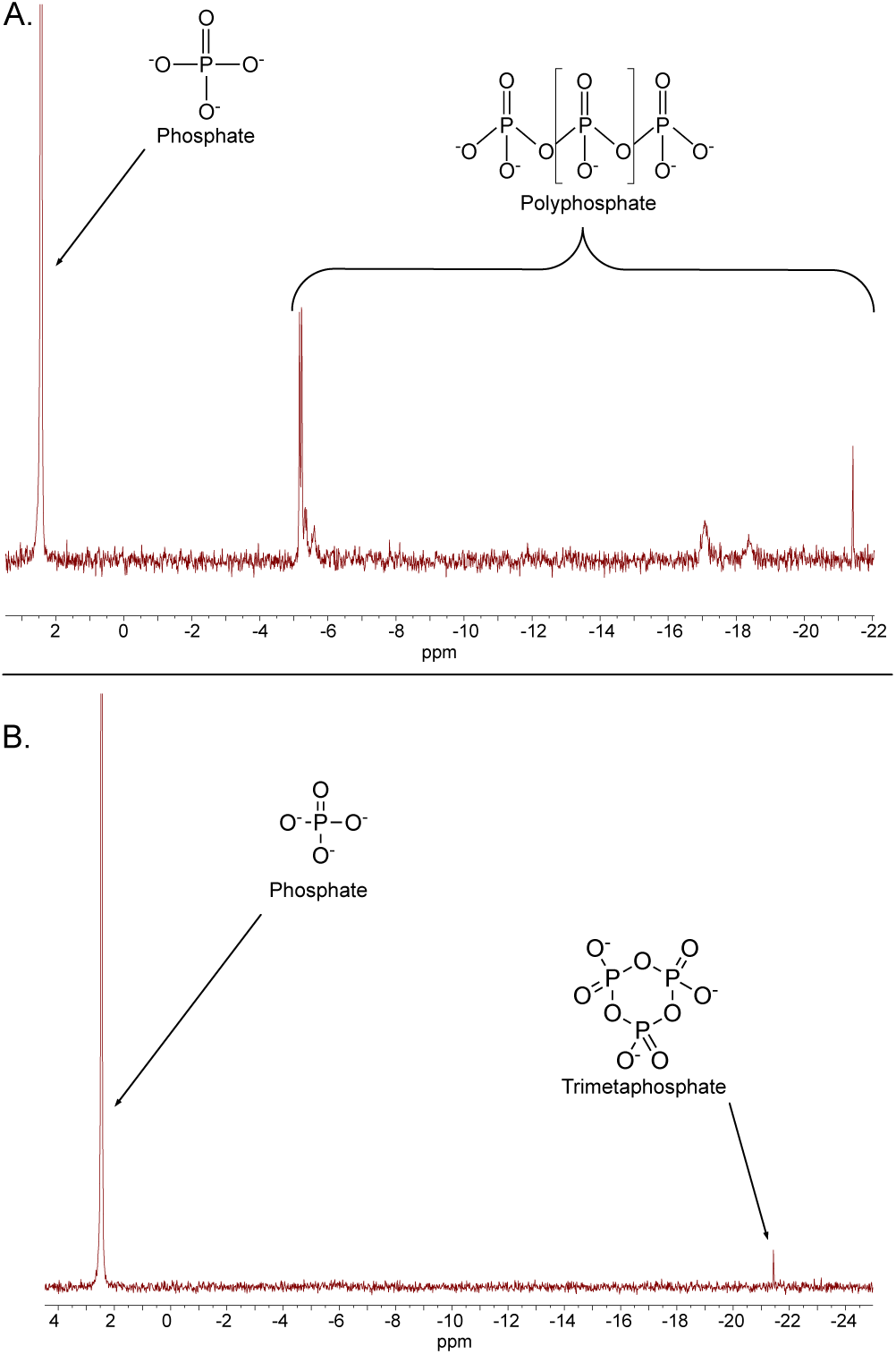
Formation of linear and cyclic condensed phosphates by phosphite oxidation over Pd/C. **(A)** ^31^P NMR spectrum of a reaction containing 200 mM phosphite and 1.5 mmol Pd/C at 80 °C for 18 h under 5 bar Ar at pH 9, reaction volume = 1.5 ml, showing the formation of linear polyphosphates in addition to phosphate. **(B)** ^31^P NMR spectrum of a parallel reaction performed under 50 °C at 18 h reveals the formation of trimetaphosphate, a cyclic triphosphate species, alongside residual phosphate.

Phosphite-dependent phosphorylations over Pd generate H_2_. Does H_2_ inhibit the phosphite oxidation reaction? We performed the oxidation reaction in the presence of 10 bar H_2_ instead of 5 bar Ar (**Fig. S10)**. The result shows that the reaction still proceeds, but with 62% lower yield (38% instead of 100% conversion). High partial pressure of H_2_ thus slows the phosphite oxidation reaction but does not inhibit it completely. During the present work we observed that reactor headspace pressure increased by amounts corresponding to stoichiometric H_2_ formation during phosphate formation according to H_2_O + HPO_3_^2–^ ® HPO_4_^2–^ + H_2_ with Δ*G*_0_′ = –46 kJ·mol^−1^ [60].

In modern serpentinizing hydrothermal systems, Pd^0^ typically occurs as Pd-awaruite [57–59], while our present reactions are catalyzed by Pd^0^ supported on carbon. Therefore, we synthesized Pd-awaruite as 5% w.t Pd in Ni_3_Fe and tested its ability to phosphorylate AMP. This Pd-awaruite catalyst was obtained through the one-step calcination/reduction of nickel, iron, and palladium nitrate precursors under H_2_/Ar, ensuring the formation of the target alloy phase. The X-ray diffraction (**Fig. S11 A**) confirmed formation of a highly crystalline alloy phase with face-centered cubic (fcc) structure, similar to Ni_3_Fe phase (JCPDS no. 88-1715). Sharp reflections indicate the formation of larger crystallite sizes, while no additional PdO-related phases or other phase impurities were detected. Further, scanning electron microscopy coupled with energy-dispersive spectroscopy (SEM-EDS) revealed a homogeneous distribution of Ni, Fe, and Pd, with a composition close to the intended formulation (wt%: Ni = 66%, Fe = 29%, Pd = 5%) (**Fig. S11 B**). SEM imaging also confirms the presence of large sintered particles with intraparticle porosity.

The catalytic results are shown in **Fig. 6**. 5% Pd- Ni_3_Fe is effective as solid-state catalyst, indicating that the most common form of Pd identified so far in serpentinizing systems, Pd-awaruite, also catalyzes phosphorylation reactions via HPO_3_^2–^ oxidation.

**Figure 6.**
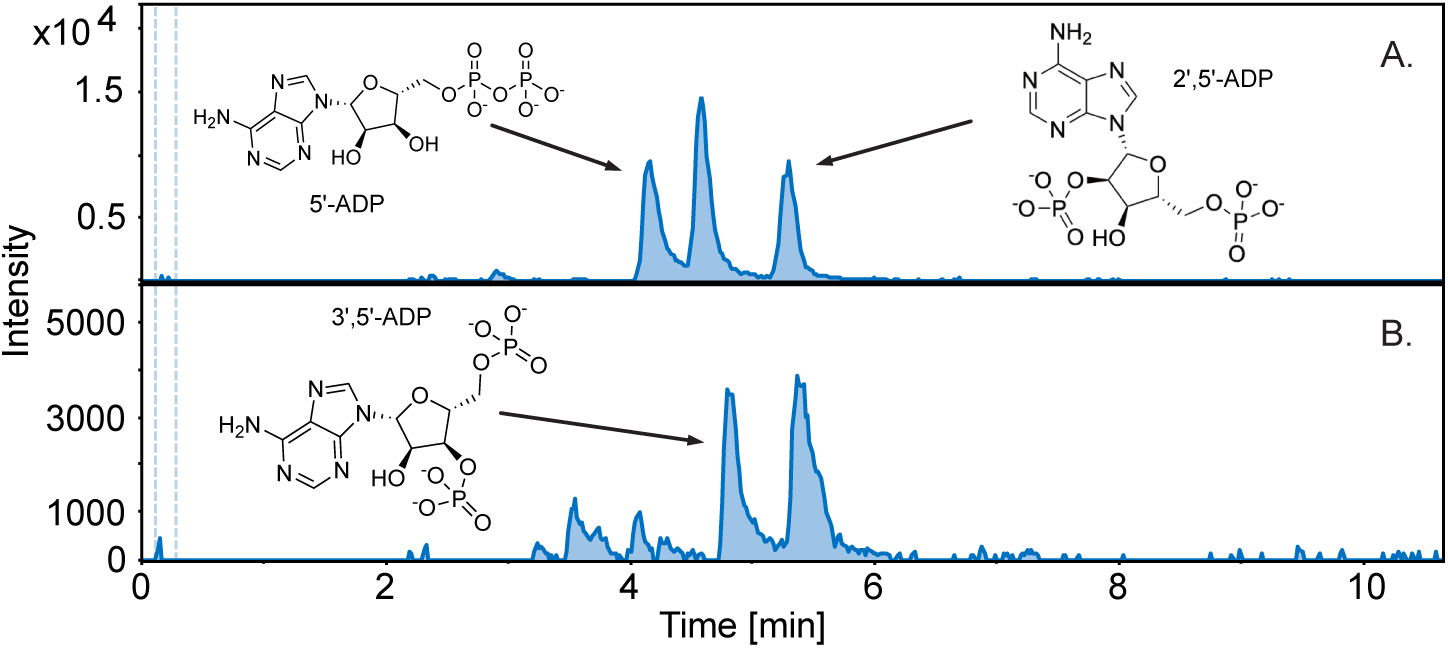
Formation of ADP via phosphite-dependent phosphorylation over Pd–Ni_3_Fe catalysts under varying conditions. (A, **B)** All experiments were performed with 100 mM AMP, 200 mM phosphite, a reaction time of 96 h, and 1.5 mmol catalyst (**Fig. S12**). (**A)** Reaction with 5% Pd–19% Ni_3_Fe at 50 °C, showing the formation of ADP from AMP with multiple peaks corresponding to different phosphorylation sites. **(B)** Reaction performed at 100 °C with 5% Pd–19% Ni_3_Fe. The different ADP isomers are depicted in their protonated forms. The yield of ADP (µM) for all three isomers combined is **A:** 13.7 and **B:** 3.8, with relative proportions of each isomer given in **Table S3**.

#### Possible mechanism

What is the phosphorylating agent? We have no direct evidence for the nature of reaction intermediates in these phosphorylations, but the products we observe are consistent with a proposal for the course of phosphite-dependent phosphorylation over Pd as outlined in **Fig. 7**. The proposal involves solid state transition metal (native palladium) catalysis as the phosphite-activating step and thus differs from the acyl phosphite-dependent mechanism proposed by Buckel [34].

**Figure 7.**
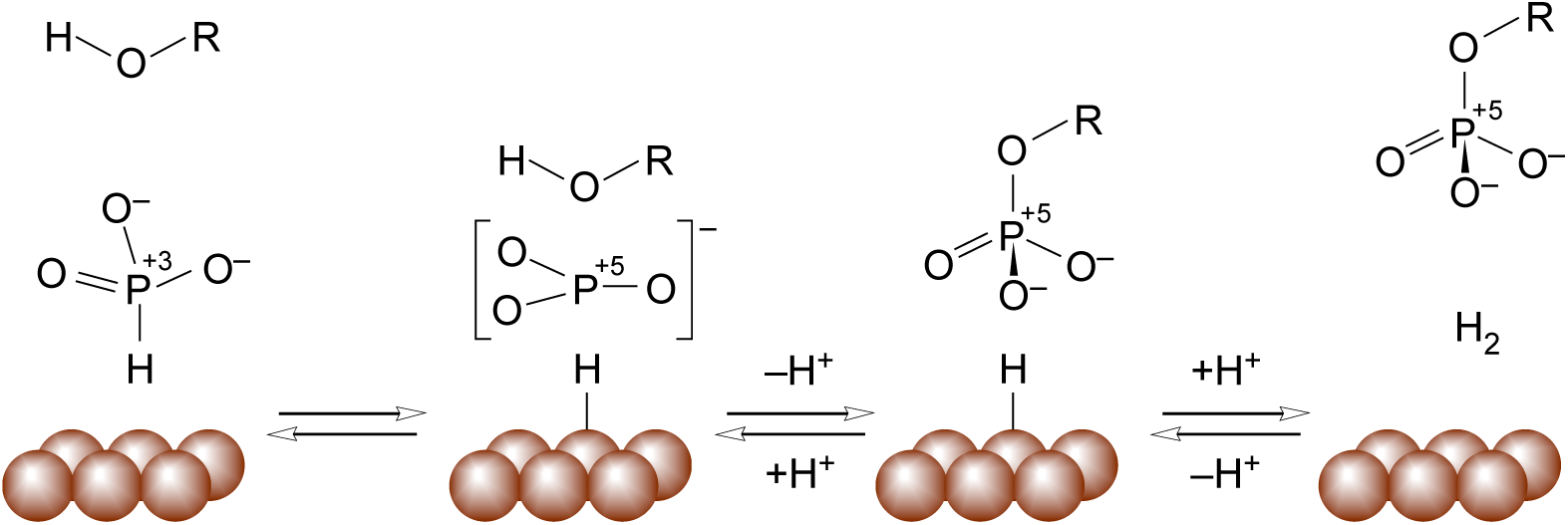
Proposed mechanism for phosphite-dependent phosphorylation on metal surfaces. A schematic proposal for the course of phosphite-dependent phosphorylation: Oxidative hydride removal, intermediate formation (metaphosphate shown as a possibility), P–O bond formation (P–N bond formation in the case of phosphocreatine), H_2_ formation.

The proposed mechanism is based in part on the proposed mechanism for *Phosphitispora fastidiosa* AdpA [33] (hydride removal) and the dissociative mechanism proposed for *Bacillus* sp. phosphite oxidase [85], but also in part on the observations (1) that implicate metaphosphate in acyl phosphate [86] and enzymatic phosphoester hydrolysis [87], (2) that some phosphorylating and dephosphorylating enzymes have been shown to generate a planar metaphosphate intermediate identifiable in crystal structures of fructose-1,6-bisphosphatase [88], in quantum mechanical studies of phosphoserine phosphatase [89] and in mechanistic studies of phosphoglucomutase [90]. The accumulation of trimetaphosphate and polyphosphates in our experiments (**Fig. 5**) would be compatible with a highly reactive [91] metaphosphate intermediate. The scheme proposed in **Fig. 7** should, in principle, be reversible, but the equilibrium lies on the side of phosphate in the presence of water (R=H), with low concentrations of phosphorylated compounds (R=alkyl, phosphoryl, acyl).

## Discussion

One general solution to the “phosphate problem” is the suggestion from origins research [92], microbial physiology [31, 34, 39], geochemistry [3, 35, 37, 41], environmental genomics [38, 40] and planetary science [93] that reduced forms of phosphorus were the source of phosphorylation at origins. Yet biochemically meaningful, physiologically compatible, and geochemically realistic phosphorylation reactions using phosphite or other reduced phosphorus compound have been lacking. We report such reactions, though we are not the first to describe non-enzymatic phosphorylation with phosphite.

Using phosphite under wet-dry cycles in the presence of air (21% O_2_) and with the addition of urea, Gull *et al.* [94] reported phosphorylation and phosphorylation of hydroxyl groups on organic compounds. Among their products they reported glycerol 1-phosphate, glycerol 2-phosphate, glycerol 1-phosphite, glycerol 2-phosphite, 5’-adenosine phosphite, 5’-AMP, 2’ and 3’-adenosine phosphites, as well as 2’- and 3’-AMP in addition to 2’-, 3’- and 5’- uridine phosphites. Among their inorganic reaction products they observed phosphate, pyrophosphate, pyrophosphite, and isohypophosphate. The drying method for phosphite with urea as a condensing agent, as is traditionally used for phosphate activation [25], yields mainly organic phosphites [94]. Instead of condensing agents, we used a transition metal catalyst, one that naturally exists at serpentinizing hydrothermal vents where phosphite is formed. We observed phosphorylated organic products and phosphorylated inorganic products, indicating that phosphite is oxidized on the catalyst surface prior to reaction with phosphorylation substrates (or water), in line with our proposal for transient metaphosphate formation during the reaction as in **Fig. 7**.

Our report is also not the first to demonstrate acetyl phosphate formation in a prebiotic context. Lohrmann and Orgel [22] obtained acetyl phosphate from reactions of phosphate with ketenes, highly reactive compounds with the general structure R_2_C=C=O, under non-physiological conditions that, like ketenes themselves, are unlikely to have existed in life-giving quantities on an early Earth covered with rocks and water [65]. Whicher *et al.* [95] also obtained acetyl phosphate, but from reactions of thioacetate with phosphate, whereby (i) thioacetate itself is typically synthesized from the reaction of H_2_S with organic anhydrides [96] and (ii) a convincing prebiotic synthesis of thioacetate is yet to be reported. More recently, Chen *et al.* [97] reported aqueous phosphorylation of acetate using diamidophosphate with sodium nitrite in a reaction driven by N_2_ release, whereby diamidophosphate is a strong, but non-physiological phosphorylating agent [98]. Pasek *et al.* [71] reported the phosphorylation of acetate using schreibersite, (Fe,Ni)_3_P, a reactive and corrosive phosphorous species that occurs in meteorites but that is not known in terrestrial chemistry. Schreibersite generates radicals and the reaction products reported [71] included non-physiological compounds resulting from a radical based (one-electron) mechanism that attacked the methyl group of acetate, rather than the carboxyl. In microbes, phosphite oxidation is NAD^+^-dependent [33], hence a two-electron, hydride transfer reaction. Our reaction products (**Fig. 1–6**) are well-known phosphorylated compounds from metabolism, our proposed reaction scheme (**Fig. 7**) involves hydride transfer.

A summary of our present findings (**Table 2)**, reveals that a single phosphorylation system in water, Pd^0^/HPO_3_^2–^, can phosphorylate a broad spectrum of biologically relevant substrates at their physiological positions. Notably, these reactions ***conserve a portion of the energy*** in the phosphite oxidation reaction (Δ*G*_0_′ = –46 kJ·mol^−1^), either as bonds with sufficient group transfer potential to phosphorylate ADP (for example acetyl phosphate or phosphocreatine), or as bonds with sufficient free energy of hydrolysis to promote further reaction. One might interpret **Table 2** as evidence that Pd/HPO_3_^2–^ generates excellent phosphorylating agents, such as ADP, phosphocreatine or acetyl phosphate, but our interpretation is more direct: Pd/HPO_3_^2–^***is*** the phosphorylating agent. In the most direct interpretation, Pd/HPO_3_^2–^ represents the primordial source of phosphorylation and phosphate as the energy currency of life [1], present from the very beginning of prebiotic chemistry, long before the origin of enzymes, substrate level phosphorylation (SLP), ATP, or gradient harnessing ATP synthases [9]. Once early evolving chemical systems and LUCA [15, 99] had invented the basic enzymatic machinery of bioenergetics, phosphorylation via HPO_3_^2–^ and solid state metals was replaced by energy metabolism in its modern state: ion pumping via redox reactions coupled to the plasma membrane with ATP synthesis via rotor stator ATP synthases and enzymatic SLP.

**Table 2.** Phosphite-dependent phosphorylations using Pd/HPO_3_^2^– in water.

| Reactant | Product | Time<br>[h] | Temp<br>[°C] | Yield<br>[%] | $\Delta G_0'$ hydr.<br>[kJ·mol<sup>-1</sup>]<sup>a</sup> |
| --- | --- | --- | --- | --- | --- |
| Acetate | Acetyl-P | 18 | 25 | 8 | -43 |
| Creatine | Phosphocreatine | 18 | 60 | 0.002 | -43 |
| AMP | ADP | 72 | 50 | 7.3 | –31 |
| AMP<sup>c</sup> | ADP | 72 | 50 | 0.013 | –31 |
| Phosphate | Trimetaphosphate | 18 | 80 | ~1<sup>d</sup> | –29<sup>b</sup> |
| Ribose | Ribose 5-P | 72 | 30 | 0.6 | –23 |
| Glucose | Glucose 6-P | 18 | 40 | 1.1 | –21 |
| Phosphate | Pyrophosphate | 18 | 80 | ~1<sup>d</sup> | –21 |
| Cytidine | Cytidine 5'-P | 72 | 30 | 0.37 | –14 |
| Serine | Phosphoserine | 18 | 50 | 49.5 | –12 |
| Glycerol | Glycerol-P | 72 | 50 | 1.7 | –9 |

Notes: <sup>a)</sup> Free energy of hydrolysis of the respective organophosphate bond in the reaction product, values from this work and the literature. For $\text{HPO}_3^{2-} + \text{H}_2\text{O} \rightarrow \text{HPO}_4^{2-} + \text{H}_2$ , $\Delta G_0' = -46.3 \text{ kJ}\cdot\text{mol}^{-1}$ [60]. <sup>b)</sup> Meyerhoff *et al.* [100] reported the free energy of hydrolysis of trimetaphosphate to three orthophosphate molecules as $-88 \text{ kJ}\cdot\text{mol}^{-1}$ or $-29.3 \text{ kJ}\cdot\text{mol}^{-1}$ per metaphosphate bond. If several reactions for the same substrate were performed, the conditions with the highest yield are given. <sup>c)</sup> This reaction was performed using 5%Pd in $\text{Ni}_3\text{Fe}$ (**Fig. 6**), all other reactions in the table refer to Pd/C catalyst. <sup>d)</sup> Small specific peak in $^{31}\text{P}$ NMR, detection limit ca. 1% conversion.

Though our yields are typically in the low % range, the process we are modelling here is a serpentinizing hydrothermal vent, a (natural) continuous flow reactor system, in which organics are continuously synthesized from H_2_, CO_2_ and NH_3_ (for a recent overview, see [56]). In the model, environmental phosphite is continuously supplied at steady state concentrations, and is activated (oxidized) for phosphorylation reactions catalyzed by immobile solid state catalysts, continuous supply over time counterbalancing modest batch yields in an origins context.

Without activation, phosphite is generally inert despite a large free energy of hydrolysis (**Table 1**) and a very negative midpoint potential (–690 mV [31]) and Δ*G*0′ (–46.3 kJ·mol^−1^ [60]) in the phosphite to phosphate conversion. Herschy *et al.* [35] estimated that phosphite, once formed, can resist environmental oxidation for thousands to millions of years, underscoring the need for catalysts to accelerate phosphite activation. The catalyst used here, native palladium, oxidizes phosphite to phosphate in water in hours (**Fig. 1–6**) as opposed to millennia [35] or seconds for the NADH-forming enzymatic reaction catalyzed by AdpA [33]. Notably, Yang and Metcalf [60] reported that alkaline phosphatase from *Escherichia coli* rapidly oxidizes phosphite to phosphate in a H_2_-producing reaction. An intermediate in the Pd-catalyzed reaction reported here forms phosphate bonds with oxygen and nitrogen atoms (**Fig. 7**); it is not known whether alkaline phosphatase [60] will phosphorylate organic substrates with phosphite. Nagaosa and Aoyama [101] reported the oxidation of phosphite to phosphate in aqueous solutions using Pd^0^ and Pd/C in the presence of air (21% O_2_). In our reactions without organics, the only available electron acceptors are protons in water, which form H_2_. In the presence of organics, many residues other than H^+^ could have become reduced during the reaction, because the midpoint potential of the phosphite to phosphate reaction is more negative than even the strongest reductants (*E*_0_ = –622 mV) produced by non-photosynthetic organisms [102]. However, we did not observe the reduction of organics, suggesting that the anaerobic oxidation reaction specifically generates metal-bound hydride and then H_2_, rather than other reduction products, suggesting in turn that an oxidized intermediate is first generated during the phosphite oxidation reaction that subsequently phosphorylates available substrates (**Fig. 7**), rather than phosphite reducing them.

Several theories about biochemical evolution posit a central role for sulfur in primordial bioenergetics: the thioester world [103] and the iron-sulfur world [8]. In de Duve’s theory [103], thioesters preceded phosphorylated compounds as the universal energy currency. In Wächtershäuser’s theory, pyrite (FeS_2_) formation preceded phosphate as the primordial energy currency [8]. Other theories posit primordial energetics provided by reactions of diverse organosulfur compounds [104, 105], in some formulations with no role at all for phosphate at the origin of metabolism [106]. A chemical theory posits cyanide, H_2_S and energy from UV light at the origin of metabolism [107]. Contrary to sulfur-based theories for energy at origins, the reactions reported here generate the main carriers of biochemical energy in microbial metabolism—phosphoanhydrides and acyl phosphates—overnight in water in the complete absence of sulfur. Thus, neither thioesters, organosulfides, metal sulfides, nor H_2_S are required for energy harnessing via phosphorylation in our experiments and by inference at metabolic origin. Furthermore it now seems possible, if not likely and contrary to earlier proposals by one of us [9], that thioesters (Δ*G*_0_′ = –32 kJ·mol^−1^) were generated from acyl phosphates (Δ*G*_0_′ = – 43 kJ·mol^−1^) in a prebiotic context, which would be more in line with their relative ranks among high-energy bonds (**Table 1**).

Theories that address the origin of phosphate-based energetics posit that substrate-level phosphorylation preceded chemiosmotic energy harnessing [9–11]. Our findings agree with that view. Moreover, serpentinizing hydrothermal vents are now known to generate the geochemical energy underlying both sources of high-energy phosphate bonds. It is known that serpentinization generates proton gradients of sufficient strength (1–3 pH units) and the right polarity (alkaline inside) to drive chemiosmotic ATP synthesis [9, 108, 109]. Consistent with that view, ATP synthase reconstituted into primordial-type vesicles consisting of primordial fatty acids alone can maintain a pH gradient of ∼1 pH unit and drive ATP synthesis [110]. Phosphite-dependent phosphorylations are also a source of continuous and *geochemically supplied* chemical energy (**Table 2**), provided *for free* in the currency of metabolism—phosphorylation—by the environment of serpentinization and stemming from the highly exergonic nature of the phosphite-oxidation reaction (**Table 1**). The environment needs to provide only water, Pd^0^, phosphite, and substrates, without the need for condensing agents, diamidophosphates, phosphides, recurrent drying, UV light, or sulfur. The involvement of a metaphosphate intermediate in the reaction mechanism (**Fig. 7**) would be very much in line with Westheimer’s arguments [1] in favor of a role for metaphosphate in biochemical phosphorylation reactions.

Spontaneous geochemical reactions under the aqueous conditions of serpentinizing hydrothermal vents now provide all three chemical energy currencies used in the metabolism of modern cells, continuously, and for free: (i) H_2_ as a diffusible reductant for the generation of reduced ferredoxin [55], a key energy currency in anaerobes [73, 111], (ii) harnessable chemiosmotic gradients [74] generated by the alkalinity of serpentinization effluent [9, 110], and now (iii) substrate level phosphorylation provided by phosphite and native metals naturally deposited in serpentinizing systems [46, 57, 58]. From the standpoint of phosphorylation and metabolic energy, this sets serpentinizing systems distinctly apart from all other chemical environments so far proposed as the site of life’s origin. Our findings place Westheimer’s question “Why nature chose phosphates” [1] in new light: It now seems possible that phosphates were not *selected*, they were *accepted* as the main aqueous energy currency that the environment had in continuous supply, primordial energy metabolism being the outcome.

## Material and Methods

Reactions contained 100 mM of each organic substrate and 200 mM sodium phosphite (NaH_2_PO_3_; Merck, Sigma-Aldrich) dissolved in distilled water and adjusted to a final volume of 1.5 mL. All organic substrates, product standards, and inorganic reagents were purchased from Merck, Sigma-Aldrich, unless stated otherwise. The pH was adjusted to 9.0. Palladium catalysts (Merck, Sigma-Aldrich) were weighed out outside a glovebox. Supported Pd/C (10 wt% Pd on activated carbon) was added at 159 mg per reaction, corresponding to approximately 0.15 mmol of elemental palladium. For comparison experiments, elemental palladium (Pd^0^) was added at the same molar loading (0.15 mmol Pd). Catalyst loadings are reported as mmol of elemental Pd. Reaction mixtures were transferred into 3 mL glass vials, sealed with screw caps, and punctured with a sterile needle to allow pressure equilibration while maintaining a closed reaction environment. The vials were placed in a stainless-steel pressure reactor equipped with a temperature controller (Berghof BR-300 with BTC-3000). The reactor was purged three times with argon and subsequently pressurized to 5 bar Ar (99.999%). Reactions were conducted at 25, 30, 50, 80, or 100 °C for incubation times of 2, 4, 18, or 72 h, as specified for each experiment. After completion, the reactor was depressurized, and the vials were immediately transferred to 2 mL microcentrifuge tubes. Samples were centrifuged for 20 min at 16,060 × g to separate the catalyst from the supernatant. The supernatants were collected for NMR analysis.

### Synthesis and Physicochemical Characterization of the Pd–awaruite Catalyst

Regarding the Pd-awaruite catalyst (5% Pd-Ni_3_Fe), nickel, iron and palladium nitrate precursors (Merck, Sigma-Aldrich) were mechanically ground and mixed before calcining/reducing them in a single step (20% H_2_/Ar, 100 ml/min, 1 °C/min, 550 °C, 8 h). After that, the catalyst was passivated by using 10% air/Ar mixture for 25 min. X-ray diffraction (XRD) pattern was recorded using a PANalytical Empyrean X-ray diffractometer with a Bragg-Brentano configuration, and Cu Kα radiation as the X-ray source (λ = 0.1544 nm). The instrument was operated with a voltage of 45 kV and a current of 40 mA. The diffractogram was collected in the 2θ region defined between 20° and 80°. After acquisition, X’Pert HighScore Plus software was employed for the analysis and background subtraction. In order to identify the crystalline phases, JPCDS database patterns were used. Scanning Electron Microscopy (SEM images) were obtained with a field-emission Apreo 2 Scanning Electron

Microscope from Thermo Fisher Scientific. The working distance was maintained at approximately 5–10 mm, and the equipment was operated under high-vacuum conditions to achieve high-resolution imaging. Elemental analysis and mappings were obtained using Energy-Dispersive Spectroscopy (EDS) and Aztec Software.

### NMR Analysis of Phosphorylation Products

Both ^31^P and ^1^H NMR spectra were acquired at the Center for Molecular and Structural Analytics (CeMSA@HHU, Heinrich Heine University Düsseldorf). ^31^P NMR spectroscopy was used to quantify phosphite consumption, phosphate formation, and the formation of phosphorylated organic species. Spectra were recorded in D_2_O using a Bruker spectrometer operating at 242.9 MHz for ^31^P with proton decoupling (waltz16), a spectral width of 96,153.844 Hz, 65,536 data points, and 16 scans. A relaxation delay (D1) of 2.0 s was used. Spectra were processed using exponential multiplication (LB = 1 Hz) and analyzed with MestReNova (version 14.2.0-26256). ^1^H NMR spectra were recorded on a Bruker Avance III spectrometer operating at 600 MHz. Samples were measured in an H_2_O/D_2_O mixture (6:1, v/v), with sodium 3-(trimethylsilyl)-1-propanesulfonate (DSS) as an internal chemical shift reference (methyl signal at 0.00 ppm). Spectra were acquired using a noesygppr1d pulse sequence with a relaxation delay (D1) of 1 s, a time-domain size of 98,520 points, and a spectral width of 12,315.271 Hz. Sixteen scans were accumulated per sample. Additional acquisition parameters included an acquisition time of 4.0 s and water suppression via presaturation. Spectra were processed using exponential multiplication (LB = 0.3 Hz) and integrated using Chenomx NMR Suite (version 9.0).

### Identification by ESI–LC–MS

The phosphorylated compounds: cytidine phosphate, phosphocreatine and ribose phosphate, were determined and quantified using the Dionex UltiMate 3000 UPLC system (Thermo Scientific, Germering, Germany) coupled to a maXis 4G (Bruker Daltonics, Bremen, Germany) quadrupole-time-of-flight (Q-TOF) mass spectrometer equipped with an electrospray (ESI) ion source. The separation was performed on a XSelect HSS T3 (100 Å, 2.5 μm, 3.0 x 150 mm; Waters) column at room temperature within a 20 min gradient at a flow rate of 0.3 ml/min using a binary gradient system with water/0.1% formic acid (FA) as mobile phase A and methanol/0.1% FA as mobile phase B. The following gradient was used, 0 to 3 min 2% B (v/v), 3 to 10 min linear increase to 50% B, until 13 min linear increase to 95% B, hold at 95% B until 15 min, linear decrease to 2% B until 16 min, and finally, hold at 2% B until 20 min. The samples were diluted by a factor of 10, 10 μl of sample was injected. The ESI–LC–MS and ESI–LC–MS/MS analysis was performed using a Q-TOF MS (maXis 4G; Bruker Daltonics). The instrument was operated in negative-ion mode and the operating conditions were as follows: dry gas (nitrogen): 8.0 l/min, dry heater: 200 °C, nebulizer pressure: 1.0 bar, capillary voltage: 4500 V. Data acquisition was carried out under control of the Compass HyStar software (version 6.0) (Bruker, Bremen, Germany). The different phosphorylated compounds were quantified from full-scan MS data (mass range 50-6000 m/z) using the DataAnalysis (version 6.1) software (Bruker, Bremen, Germany). The quantification is based using calibration curves obtained with commercially available standards. MS/MS spectra and the specific fragments for phosphorylated compounds in negative mode of m/z = 78.95 for PO_3_^-^ and m/z = 96.97 for H_2_PO_4_^-^ provided confirmation that it is a phosphorylated molecule.

## Abbreviations

AdpA: AMP-dependent phosphite dehydrogenase
EDS: energy-dispersive spectroscopy
ESI–LC–MS: electrospray ionization liquid chromatography–mass spectrometry
FA: formic acid
Fcc: face-centered cubic
NMR: nuclear magnetic resonance
Q-TOF: quadrupole-time-of-flight
SEM: Scanning Electron Microscopy (SEM images)
SLP: substrate-level phosphorylation
XRD: X-ray diffraction

## Author contributions

WFM, MLS, NKH, SM, JMP, HT conceived and designed the study. MLS, NKH, SM, JMP and HT carried out laboratory reactions and analytics. WFM, MLS, NKH, SM, JMP and HT designed laboratory experiments and interpreted the data. All authors contributed to results discussion. WFM, MLS and NKH drafted the manuscript. All authors edited the manuscript and approved the final version.

## Acknowledgements

We thank the **Ce**nter for **M**olecular and **S**tructural **A**nalytics, Heinrich Heine University (CeMSA@HHU) for recording the NMR-spectroscopic data. We thank Maximilian Burmeister, Eleni Dafni, Max Brabender, Joseph Moran, and Mirko Basen for many helpful discussions and Dr. Katy Evans and Stefano Tenuta for discussions on Pd and platinum group metals in serpentinizing systems. We particularly thank Bernhard Schink for helpful suggestions and the proposal to investigate palladium as a catalyst.

This project has received funding from the European Research Council (ERC) under the European Union’s Horizon 2020 research and innovation program (grant agreement no. 101018894). For funding, W.F.M. thanks the ERC (101018894), W.F.M and H.T. thank the Deutsche Forschungsgemeinschaft (DFG) (MA1426/21-1/TU315/8-1) and the Volkswagen Foundation (Grant 96_742), H.T. thanks the Spanish Ministry of Science, Innovation and Universities for an ATRAE grant.

## List of Supplementary Figures/Tables

**Figure S1.** Raw ^31^P and ^1^H NMR data for ADP formation under phosphite-driven phosphorylation conditions.

**Figure S2.** ^1^H NMR and ESI–LC–MS data supporting cytidine formation under phosphite-driven phosphorylation conditions.

**Figure S3.** ^1^H NMR and ESI–LC–MS data supporting ribose transformation under phosphite-driven phosphorylation conditions.

**Figure S4.** NMR data supporting glucose phosphorylation under phosphite-driven conditions.

**Fig. S5.** Formation of glycerol phosphate monitored by ^31^P and ^1^H NMR under phosphite-driven phosphorylation conditions.

**Figure S6.** ^31^P and ^1^H NMR spectra of serine phosphorylation under phosphite-driven conditions with Pd/C.

**Figure S7.** ^31^P and ^1^H NMR spectra of serine phosphorylation under phosphite-driven conditions with Pd^0^.

**Figure S8.** ^31^P and ^1^H NMR spectra of acetate–phosphite–Pd/C suspensions under phosphite-driven conditions.

**Figure S9.** NMR and ESI–LC–MS data supporting phosphocreatine formation under phosphite-driven conditions.

**Figure S10.** Phosphite oxidation to phosphate under hydrogen atmosphere.

**Table S1.** Raw NMR spectral data and corresponding product yields (mM).

**Table S2.** Raw ESI-LC–MS data and corresponding calculated product yields obtained from reactions containing Phi (200 mM) and ribose, cytidine, and creatine (100 mM each).

**Table S3:** Raw ESI-LC–MS data and calculated product yields obtained from reactions containing AMP (100 mM) and Phi (200 mM).

