## Supplemental figures and tables for "Hydrothermal origin of metabolic phosphorylation"

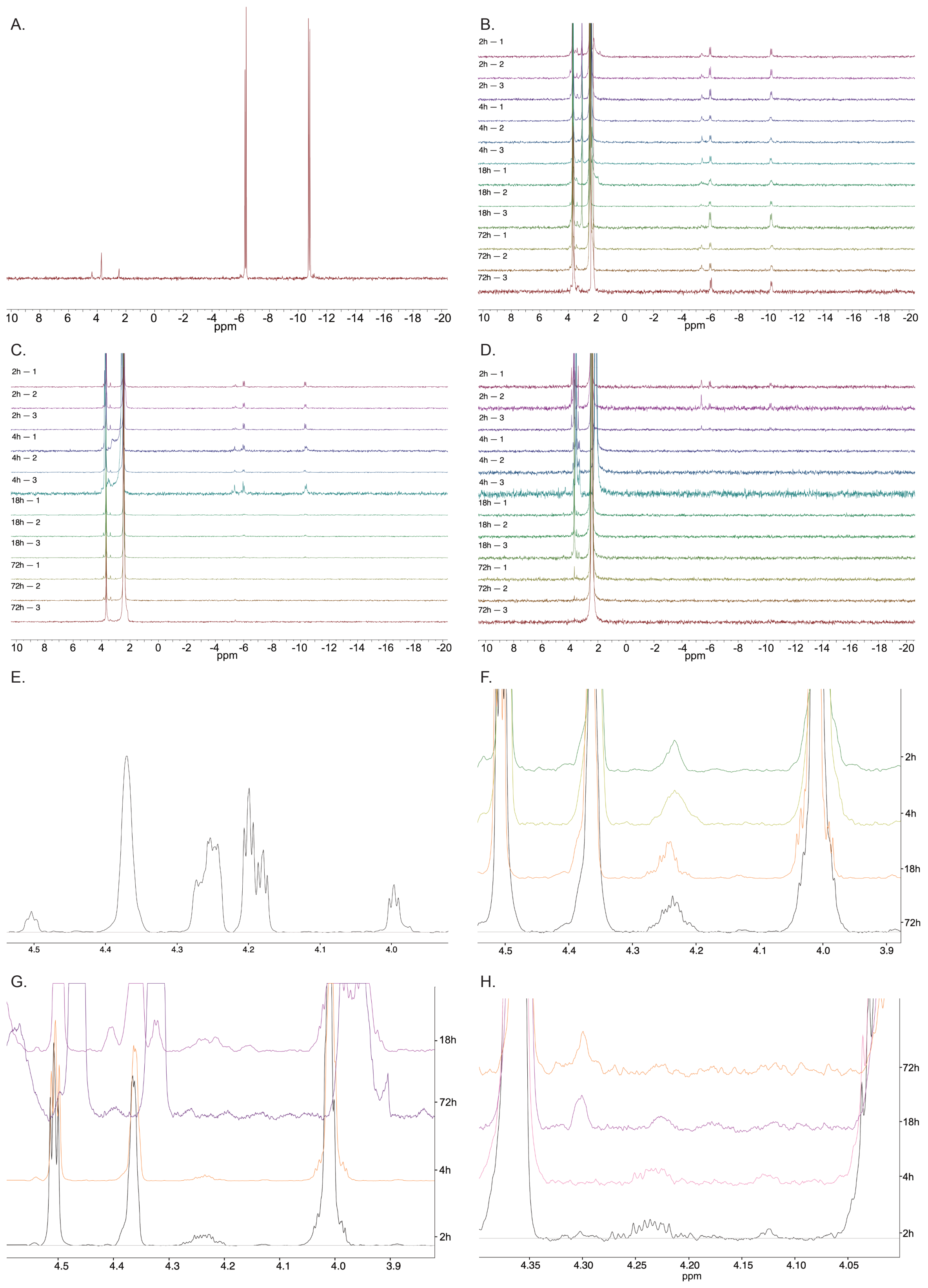

A.
B.
C.
D.
E.
F.
G.
H.
2h — 1
2h — 2
2h — 3
4h — 1
4h — 2
4h — 3
18h — 1
18h — 2
18h — 3
72h — 1
72h — 2
72h — 3
10 8 6 4 2 0 -2 -4 -6 -8 -10 -12 -14 -16 -18 -20
ppm
4.5 4.4 4.3 4.2 4.1 4.0 3.9
2h
4h
18h
72h
4.35 4.30 4.25 4.20 4.15 4.10 4.05

I.

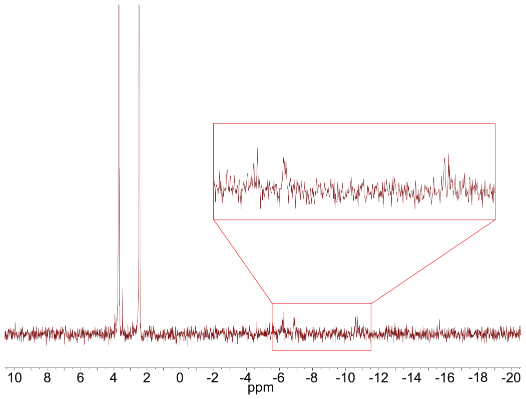

**Figure S1. Raw $^{31}P$ and $^{1}H$ NMR data for ADP formation under phosphite-driven phosphorylation conditions. (A)** $^{31}P$ NMR spectrum of an ADP standard (100 mM). **(B–D)** Raw $^{31}P$ NMR spectra corresponding to the data shown in **Fig. 1A**, recorded at 50 °C **(B),** 80 °C **(C)**, and 100 °C **(D)** for reaction times of 2 h, 4 h, 18 h, and 72 h (triplicates shown for each condition). **(E)** $^{1}H$ NMR spectrum of an ADP standard (100 mM). **(F–H)** Raw $^{1}H$ NMR spectra showing ADP formation at 50 °C **(F)**, 80 °C **(G)**, and 100 °C **(H)**, with one representative spectrum per time point (2 h, 4 h, 18 h, and 72 h). Changes in peak patterns reflect the formation and evolution of ADP over time. **(I)** $^{31}P$ NMR spectrum of a reaction mixture containing phosphite (75 mM) and AMP (100 mM), recorded after 18 h at 50 °C in the presence of 1.5 mmol Pd/C.

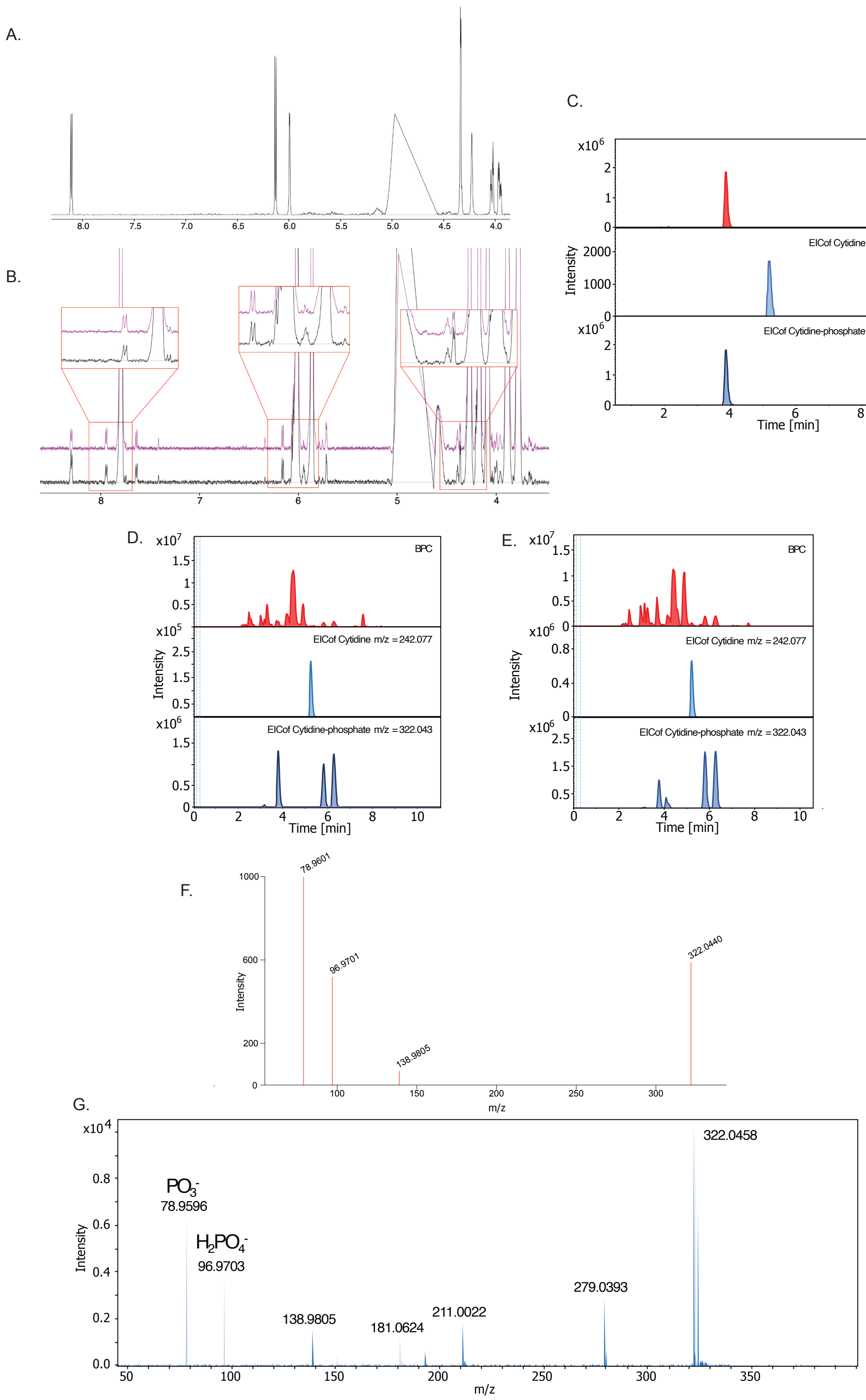

**Figure S2. ¹H NMR and ESI–LC–MS data supporting cytidine formation under phosphite-driven phosphorylation conditions. (A)** ¹H NMR spectrum of a cytidine-5′-

phosphate standard (100 mM). **(B)** $^{1}H$ NMR spectra corresponding to the data shown in **Fig. 1D**. Insets show magnified regions of key signals, demonstrating the agreement between experimental spectra and reference peaks of cytidine. **(C)** ESI–LC–MS analysis of a cytidine 5′-phosphate standard (0.01 mmol) recorded in negative ion mode, showing the base peak chromatogram (top) and extracted ion chromatograms (bottom) for cytidine and cytidine phosphate. **(D)** ESI–LC–MS analysis of the Pd/C-catalyzed reaction at 50 °C after 72 h, showing the base peak chromatogram (top) and extracted ion chromatograms (bottom) for cytidine and cytidine phosphate. Three distinct peaks are observed in the extracted ion chromatograms, consistent with phosphorylation at different positions; among these, cytidine 5′-phosphate is detected based on comparison with an authentic standard (**Figure S2 C**). **(E)** ESI–LC–MS analysis of the Pd/C-catalyzed reaction at 30 °C after 72 h, showing the base peak chromatogram (top) and extracted ion chromatograms (bottom) for cytidine and cytidine phosphate. **(F)** Reference MS/MS spectrum of cytidine-5′-phosphate obtained from MassBank (accession ID: MSBNK-RIKEN-PR100752), showing characteristic fragment ions including m/z 322.044. **(G)** Experimental MS/MS spectrum of the dominant product peak, showing fragment ions consistent with phosphorylated cytidine. Fragmentation patterns match the reference spectrum **(F)**, supporting the assignment as cytidine phosphate.

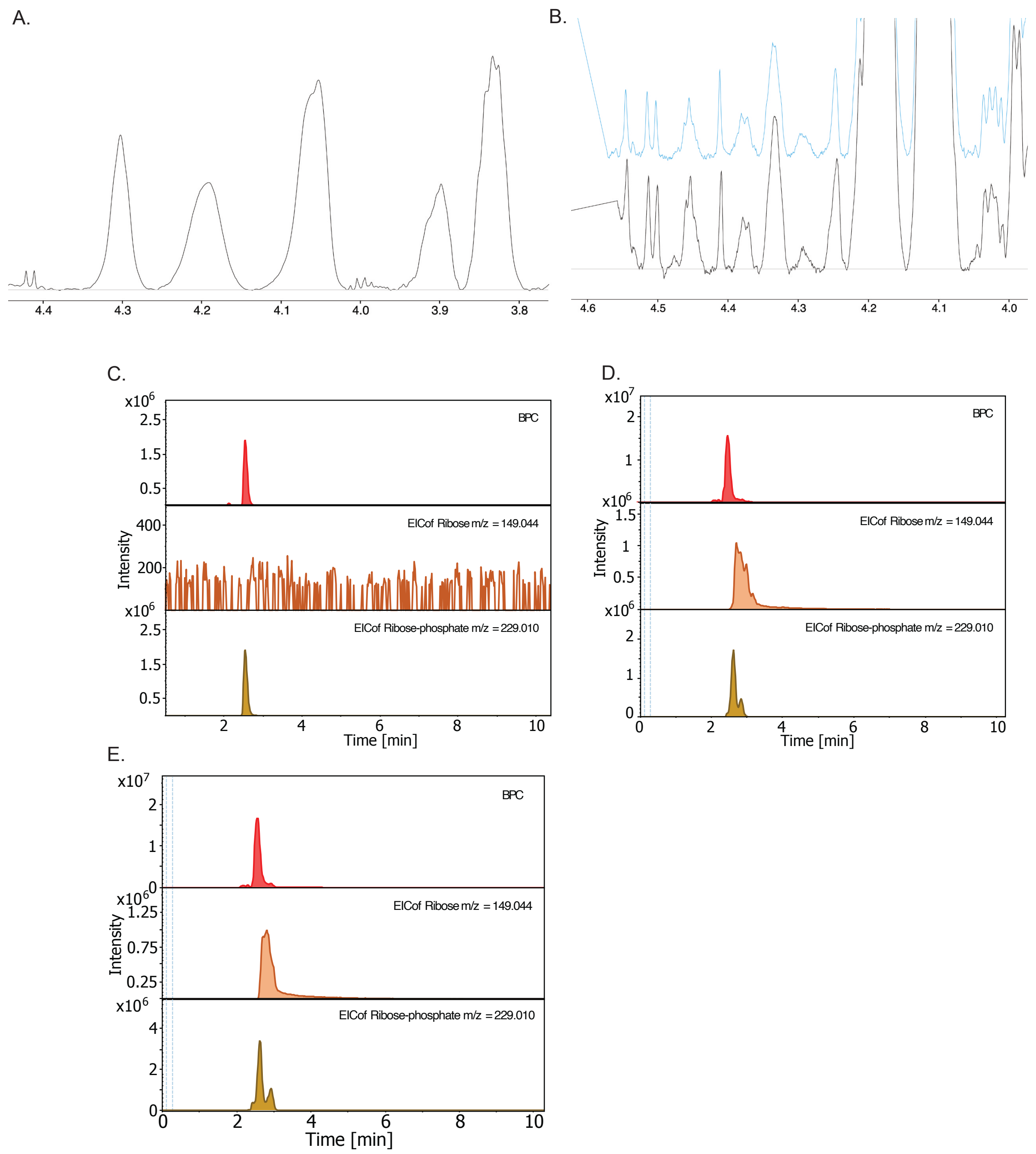

**Figure S3. [1]H NMR and ESI–LC–MS data supporting ribose transformation under phosphite-driven phosphorylation conditions. (A)** [1]H NMR spectrum of a ribose 5-phosphate standard (100 mM). **(B)** Raw [1]H NMR spectra corresponding to the data shown in **Fig. 2A**, illustrating signal evolution over time (18 h and 72 h). **(C)** ESI–LC–MS analysis of a ribose 5-phosphate standard (0.01 mmol) recorded in negative ion mode, showing the base peak chromatogram (top) and extracted ion chromatograms (bottom) for ribose and ribose phosphate. **(D)** ESI–LC–MS spectrum of the ribose reaction mixture obtained with Pd/C at 50 °C after 72 h, showing the base peak chromatogram (top) and extracted ion chromatograms (bottom) for ribose and ribose phosphate. Two distinct peaks are observed in the extracted ion chromatograms, likely reflecting phosphorylation at different positions. Comparison with an authentic ribose-5-phosphate standard (**Figure S3 C**) indicates that the most intense peak corresponds to ribose-5-phosphate. **(E)** ESI–LC–MS spectrum of the ribose reaction mixture obtained with Pd/C at 30 °C after 72 h, showing the base peak chromatogram (top) and extracted ion chromatograms (bottom) for ribose and ribose phosphate.

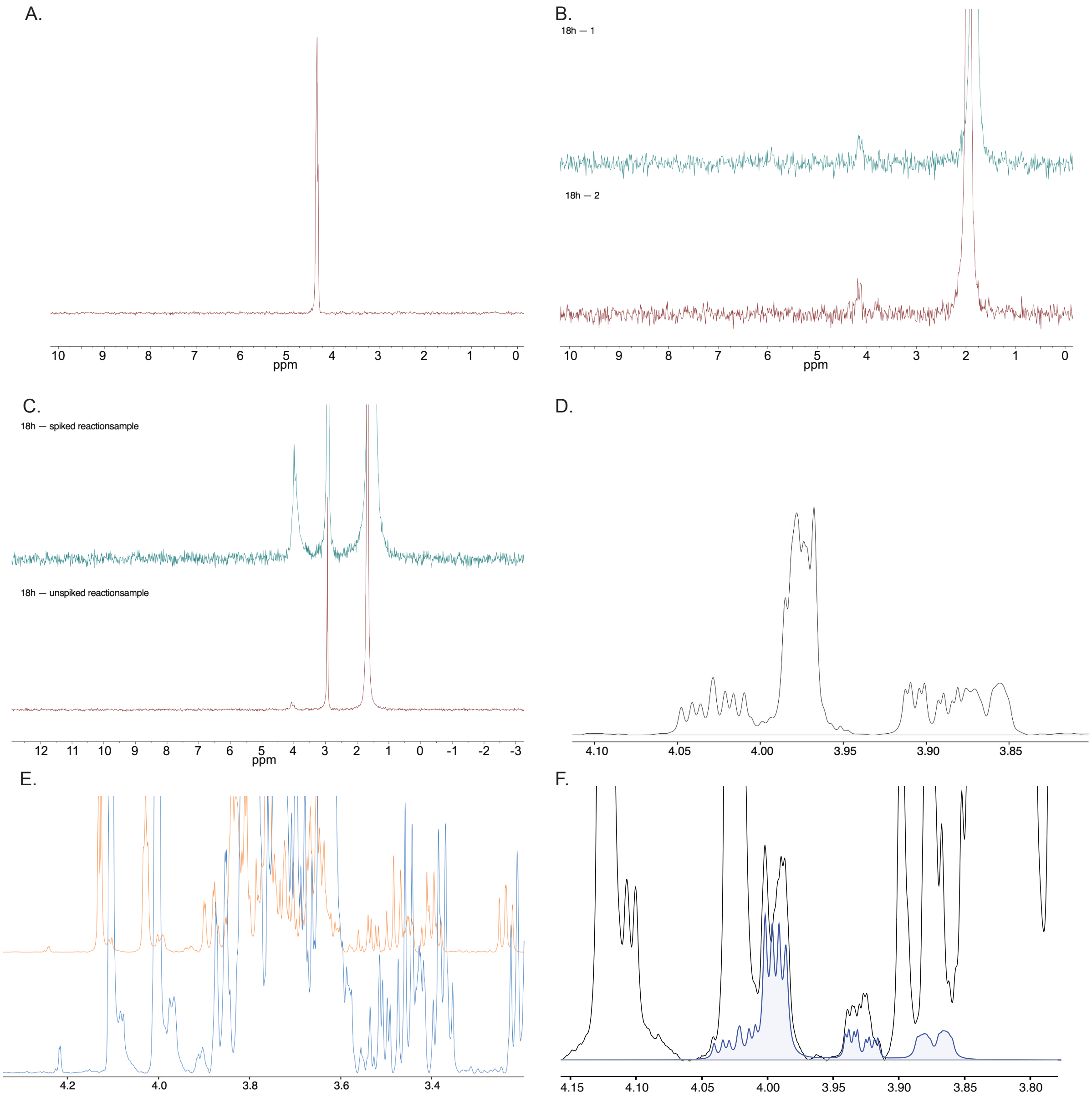

**Figure S4. NMR data supporting glucose phosphorylation under phosphite-driven conditions. (A)** $^{31}P$ NMR spectrum of a glucose 6-phosphate standard (100 mM). **(B)** Raw $^{31}P$ NMR spectrum of the reaction mixture after 18 h, corresponding to the data shown in **Fig. 2C**. **(C)** $^{31}P$ NMR spectra of the reaction mixture before (bottom) and after (top) spiking with 10 mM glucose 6-phosphate standard, confirming signal assignment. **(D)** $^{1}H$ NMR spectrum of a glucose 6-phosphate standard (100 mM). **(E)** Additional raw $^{1}H$ NMR data of the reaction mixture supporting **Fig. 2C**. **(F)** $^{1}H$ NMR spectrum of the reaction mixture with overlay of the Chenomx internal library (blue), supporting assignment of glucose-derived phosphorylation products.

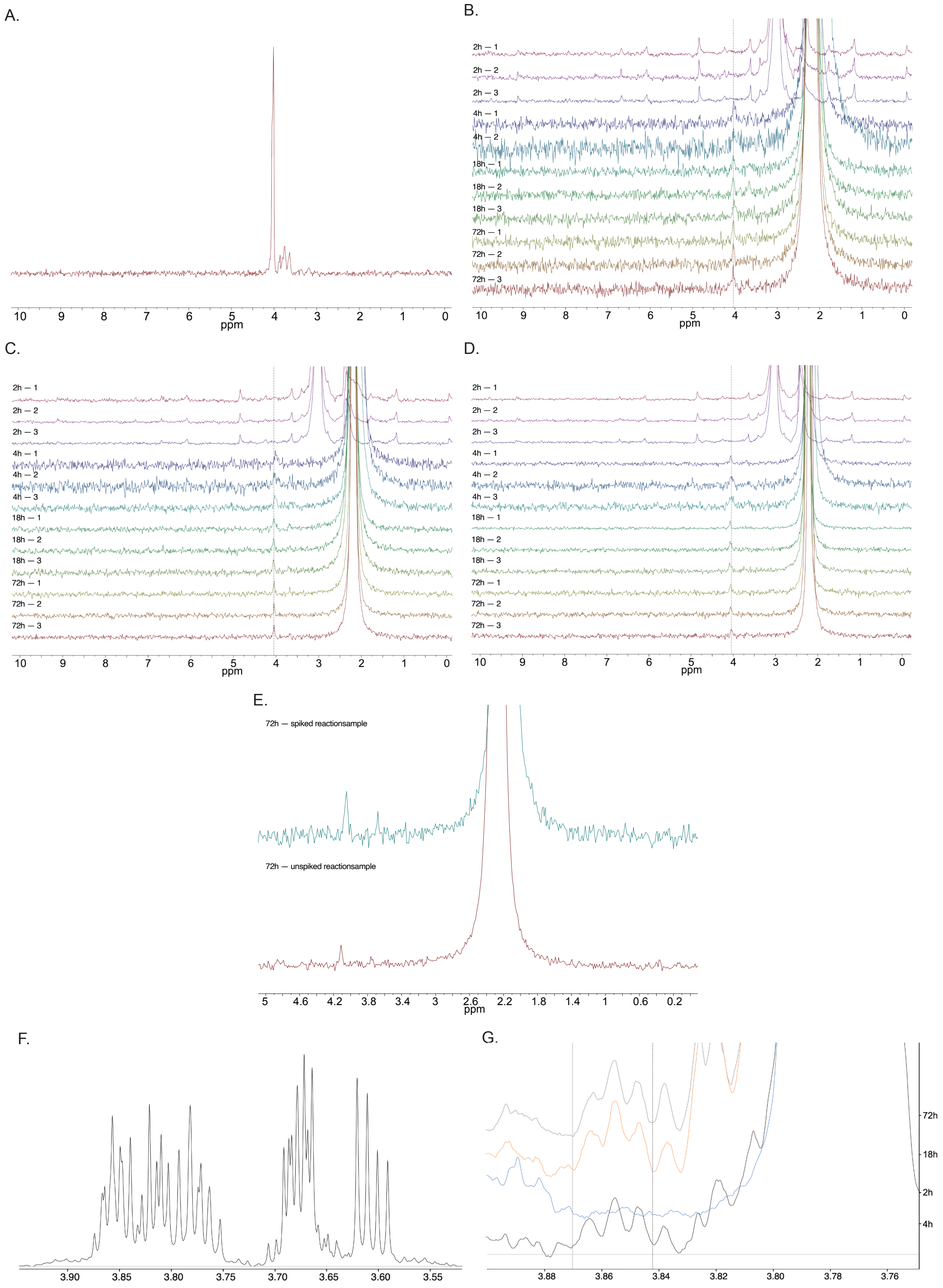
A.
B.
C.
D.
E.
F.
G.
ppm
2h — 1
2h — 2
2h — 3
4h — 1
4h — 2
4h — 3
18h — 1
18h — 2
18h — 3
72h — 1
72h — 2
72h — 3
72h — spiked reactionsample
72h — unspiked reactionsample
72h
18h
2h
4h

H.

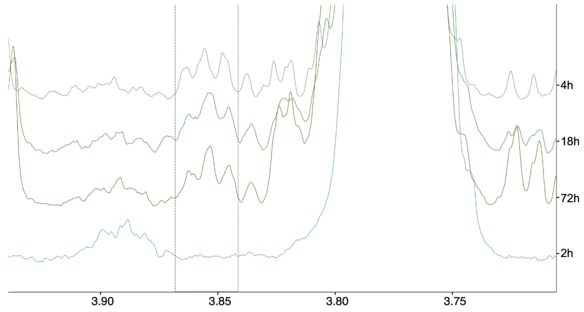

I.

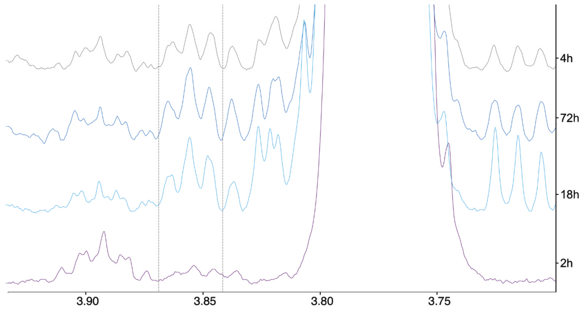

**Figure S5. Formation of glycerol phosphate monitored by $^{31}$P and $^{1}$H NMR under phosphite-driven phosphorylation conditions. (A)** $^{31}$P NMR spectrum of a glycerol 2-phosphate standard (100 mM). **(B–D)** Time-resolved $^{31}$P NMR spectra corresponding to **Fig. 3A**, recorded at 50 °C **(B)**, 80 °C **(C)**, and 100 °C **(D)**, recorded at 2 h, 4 h, 18 h, and 72 h (triplicates shown). **(E)** $^{31}$P NMR spectra of the reaction mixture before (bottom) and after (top) spiking with 10 mM glycerol 2-phosphate standard, confirming signal assignment. **(F)** $^{1}$H NMR spectrum of a glycerol 2-phosphate phosphate standard (100 mM). **(G–I)** Time-resolved $^{1}$H NMR spectra corresponding to **Fig. 3A**, recorded at 50 °C **(G)**, 80 °C **(H)**, and 100 °C **(I)**, with one representative spectrum per time point (2 h, 4 h, 18 h, and 72 h). Changes in peak patterns reflect the formation and evolution of glycerol phosphate over time.

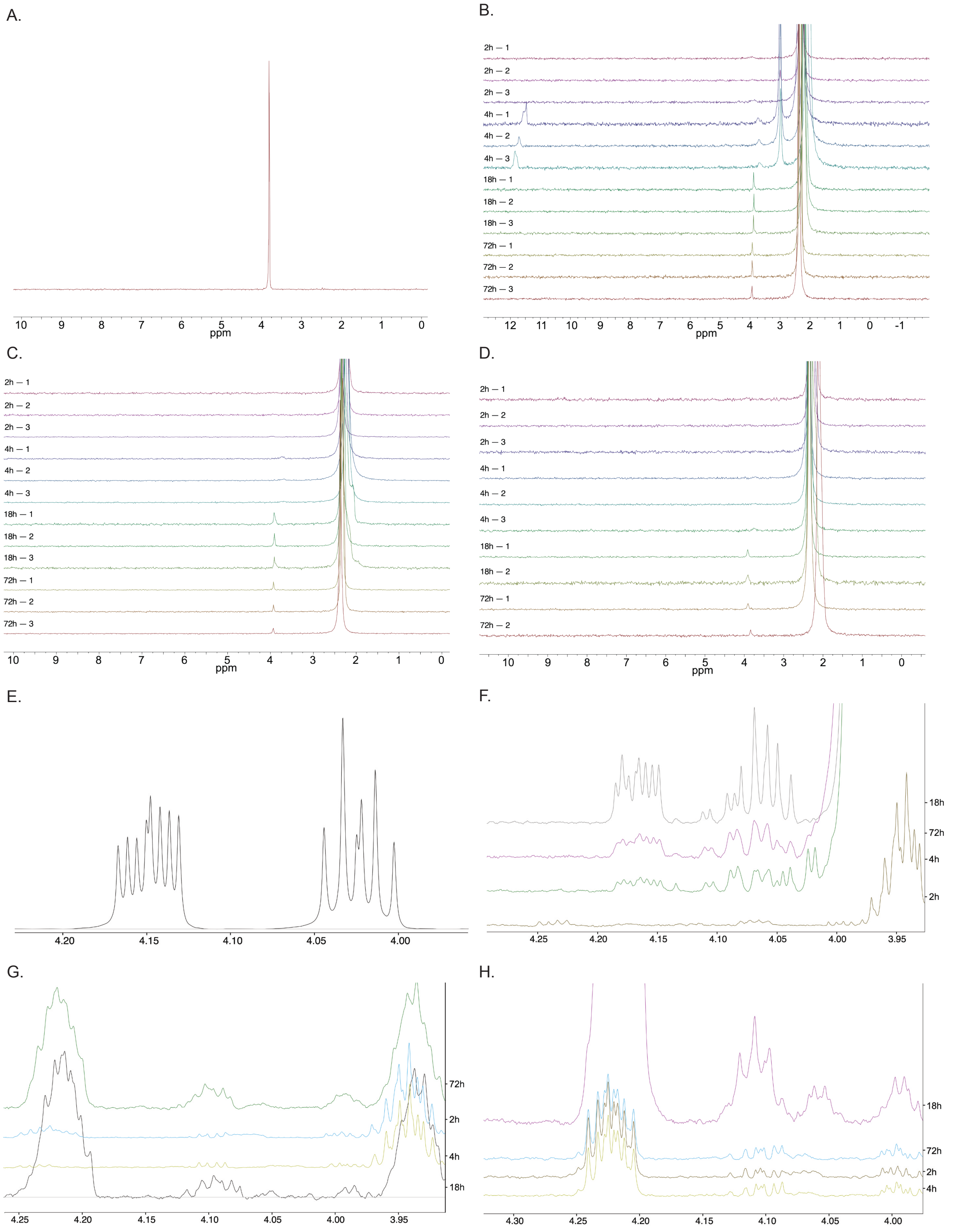

**Figure S6. $^{31}P$ and $^{1}H$ NMR spectra of serine phosphorylation under phosphite-driven conditions with Pd/C. (A)** $^{31}P$ NMR spectrum of a phosphoserine standard (100 mM). **(B–D)** Raw $^{31}P$ NMR spectra corresponding to the data shown in **Fig. 3D**, recorded for reactions at 50 °C **(B)**, 80 °C **(C)**, and 100 °C **(D)** at 2 h, 4 h, 18 h, and 72 h (duplets/triplicates shown). **(E)** $^{1}H$ NMR spectrum of a phosphoserine standard. **(F–H)** Raw $^{1}H$ NMR spectra corresponding to **Fig. 3D** under identical conditions at 50 °C **(F)**, 80 °C **(G)**, and 100 °C **(H)**, with one representative spectrum shown per time point. Spectral features are consistent with the formation of phosphorylated serine species.

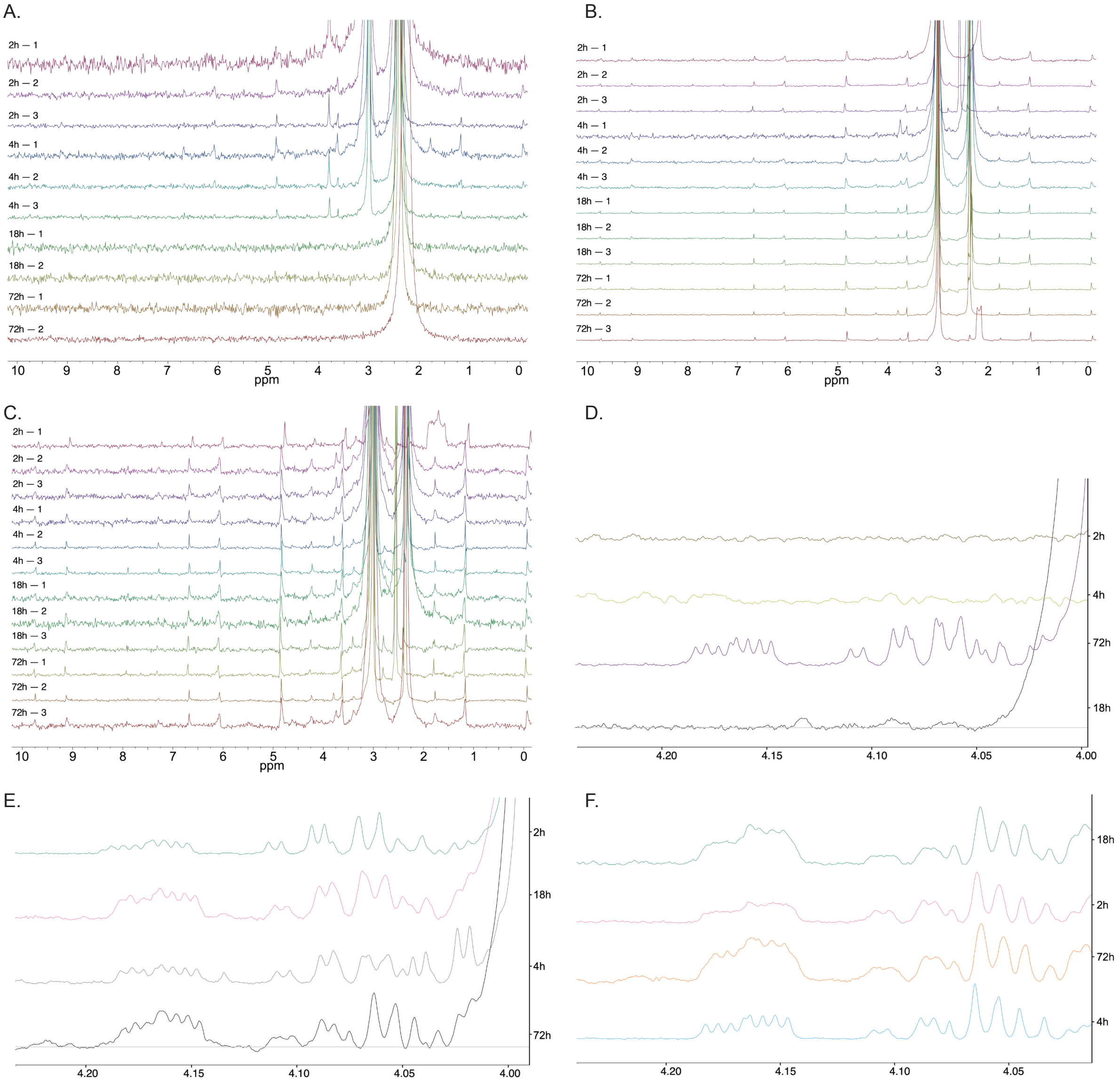

**Figure S7. $^{31}$P and $^{1}$H NMR spectra of serine phosphorylation under phosphite-driven conditions with Pd$^{0}$. (A–C)** Raw $^{31}$P NMR spectra corresponding to **Fig. 3E**, recorded for reactions at 50 °C **(A)**, 80 °C **(B)**, and 100 °C **(C)** at 2 h, 4 h, 18 h, and 72 h (triplicates shown). **(D–F)** Raw $^{1}$H NMR spectra of the corresponding reactions under identical conditions at 50 °C **(D)**, 80 °C **(E)**, and 100 °C **(F)**, with one representative spectrum shown per time point.

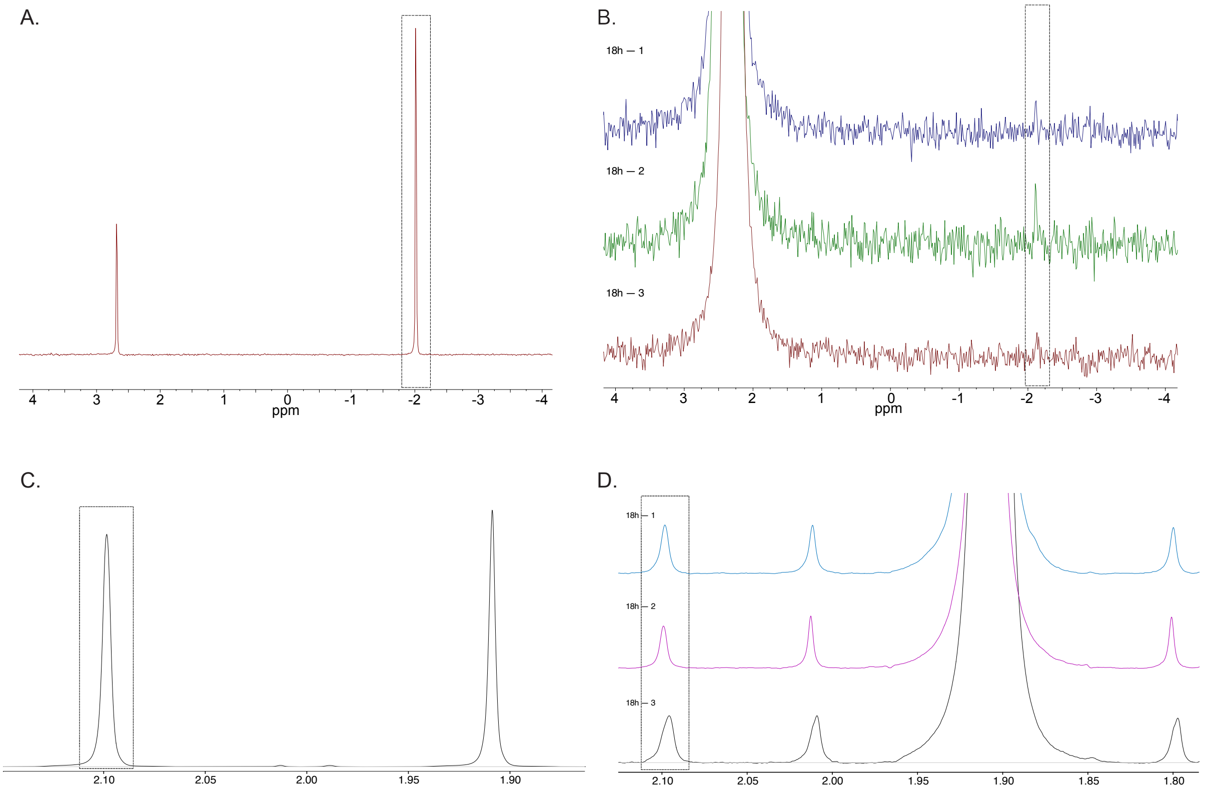

**Figure S8. $^{31}P$ and $^{1}H$ NMR spectra of acetate–phosphite–Pd/C suspensions under phosphite-driven conditions. (A)** $^{31}P$ NMR spectrum of an acetyl phosphate standard (100 mM). **(B)** Raw $^{31}P$ NMR spectra of reactions containing 200 mM phosphite, 100 mM acetate, and 1.5 mmol Pd/C in water under 5 bar argon at 25 °C. **(C)** $^{1}H$ NMR spectrum of an acetyl phosphate standard (100 mM). **(D)** Raw $^{1}H$ NMR spectra of the corresponding reactions under identical conditions.

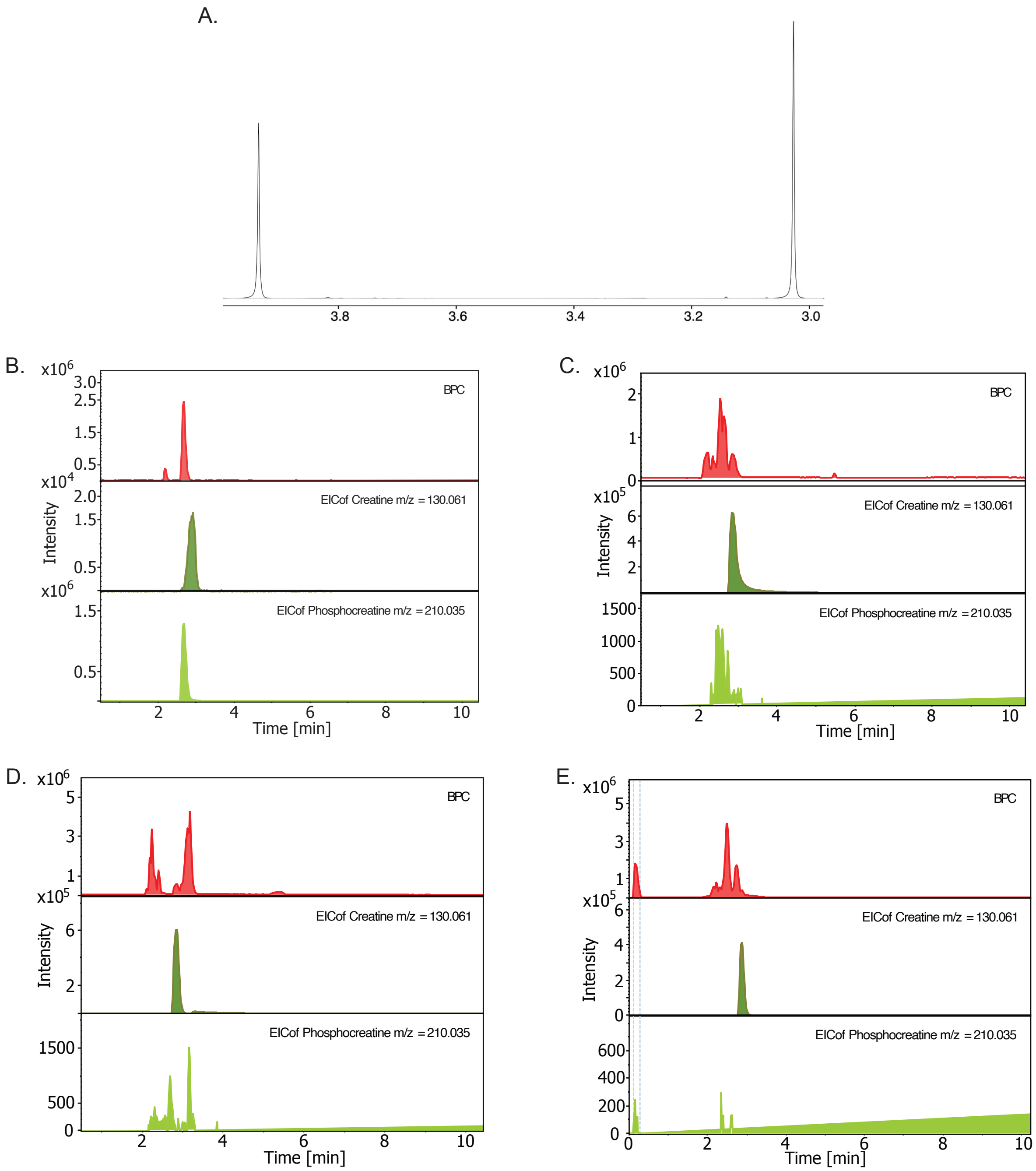

**Figure S9. NMR and ESI–LC–MS data supporting phosphocreatine formation under phosphite-driven conditions. (A)** $^{1}H$ NMR spectrum of a phosphocreatine standard. **(B)** ESI–LC–MS spectrum of a phosphocreatine standard (0.5 mmol) recorded in negative ion mode, showing the base peak chromatogram (top) and extracted ion chromatograms (bottom) for creatine and creatinephosphate. **(C)** ESI–LC–MS spectrum of the creatine reaction mixture obtained with $Pd^0$ at 60 °C after 18 h, showing the base peak chromatogram (top) and extracted ion chromatograms (bottom) for creatine and creatinephosphate. **(D)** ESI–LC–MS spectrum of the creatine reaction mixture obtained with Pd/C at 60 °C after 18 h, showing the base peak chromatogram (top) and extracted ion chromatograms (bottom) for creatine and creatinephosphate. **(E)** ESI–LC–MS spectrum of the creatine reaction mixture obtained with Pd/C at 60 °C after 72 h, showing the base peak chromatogram (top) and extracted ion chromatograms (bottom) for creatine and creatinephosphate.

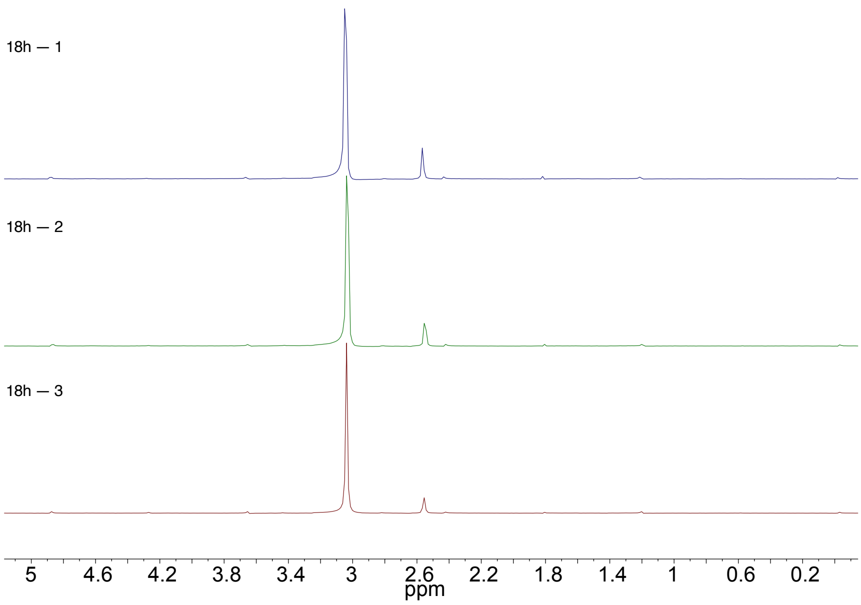

**Figure S10. Phosphite oxidation to phosphate under hydrogen atmosphere.** $^{31}$P NMR spectra of phosphite (200 mM) after reaction under 10 bar $H_2$ at 100 °C for 18 h in the presence of 1.5 mmol Pd/C. Quantification reveals the formation of 72 mM phosphate.

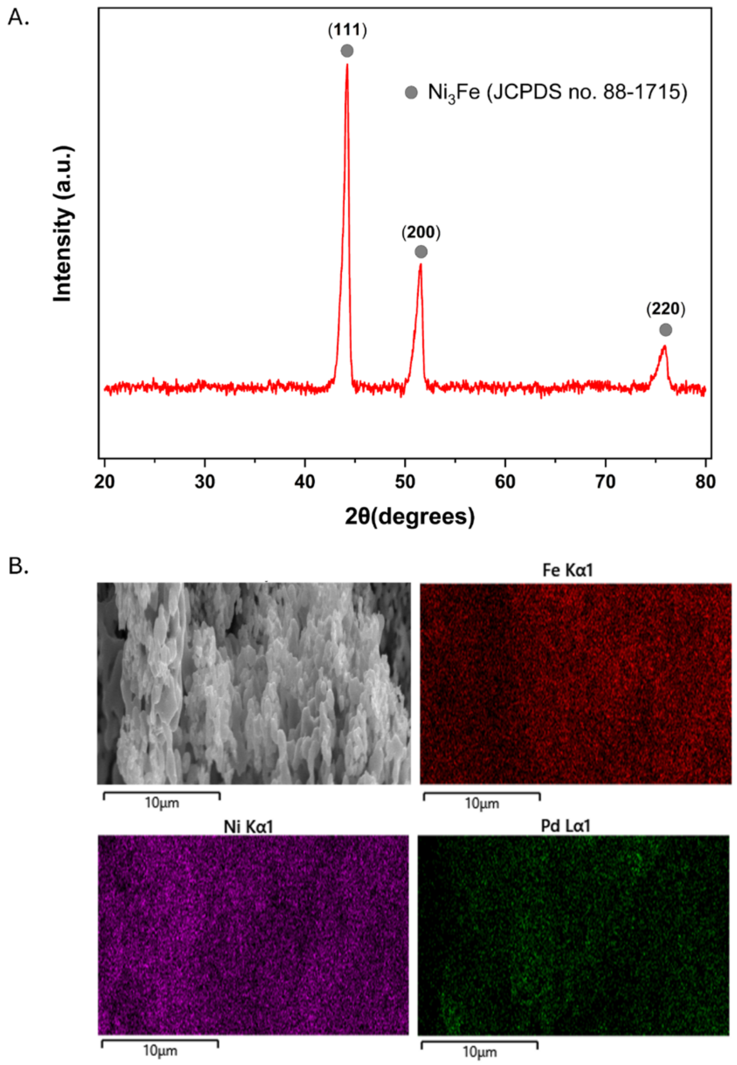

**Figure S11. Structural characterization of Pd-awaurite. (A)** X-ray diffractogram of the 5 wt.% Pd-$Ni_3Fe$ catalyst, showing the crystallographic reflections from *fcc* $Ni_3Fe$ (JCPDS no. 88-1715) **(B)** SEM images and corresponding SEM-EDS mappings showing the elemental distribution of the 5 wt.% Pd-$Ni_3Fe$ catalyst.

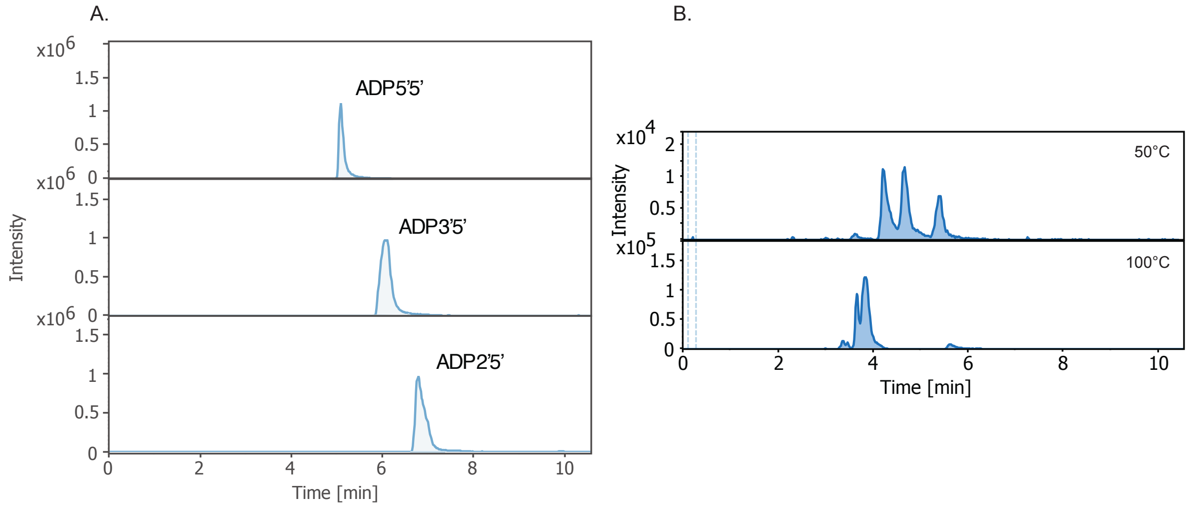

**Figure S12. ADP formation under $H_2$-driven phosphorylation conditions. (A)** ESI–LC–MS spectra of ADP standards (5′,5′-, 3′,5′-, and 2′,5′-linked isomers). **(B)** ESI–LC–MS chromatograms of reactions conducted with catalysts (1% Pd-19% $Ni_3Fe$ and 5% Pd-$Ni_3Fe$, 0.015 mmol) at 50 or 100 °C for 18 h, showing product formation under different catalyst compositions.

**Table S1:** Raw NMR spectral data and corresponding product yields (mM).

Adenosine diphosphate, Pd/C

| | 2 h | | | 4 h | | | 18 h | | | 72 h | | |
|---|---|---|---|---|---|---|---|---|---|---|---|---|
| 50°C | 4.27 | 4.24 | | 4.25 | 4.24 | 4.25 | 5.94 | 5.84 | 5.93 | 7.57 | 7.25 | 7.16 |
| 80°C | 2.89 | 2.46 | 3.02 | 1.41 | 2.17 | | 0.91 | 1.03 | 0.92 | n.d. | n.d. | n.d. |
| 100°C | 0.56 | 0.36 | 0.44 | 0.26 | 0.34 | | 0.10 | 0.11 | | n.d. | n.d. | n.d. |

Phosphoserine, Pd/C

| | 2 h | | | 4 h | | | 18 h | | | 72 h | | |
|---|---|---|---|---|---|---|---|---|---|---|---|---|
| 50°C | 0.25 | 0.39 | 0.52 | 1.43 | 1.78 | 1.99 | 50.70 | 48.29 | | 13.45 | 13.93 | 15.00 |
| 80°C | 0.53 | 0.31 | 0.65 | 1.50 | 1.42 | 1.75 | 27.67 | 25.74 | 29.72 | 13.55 | 12.23 | 14.02 |
| 100°C | | 1.65 | 1.82 | 1.05 | 1.19 | 1.30 | 6.58 | 5.65 | 5.49 | 2.33 | 2.91 | |

Phosphoserine, Pd/0

| | 2 h | | | 4 h | | | 18 h | | | 72 h | | |
|---|---|---|---|---|---|---|---|---|---|---|---|---|
| 50°C | n.d. | n.d. | n.d. | n.d. | n.d. | n.d. | n.d. | n.d. | n.d. | 0.20 | 0.16 | |
| 80°C | 0.25 | 0.12 | | 0.35 | 0.40 | | 0.71 | 0.47 | | 0.91 | 0.59 | 0.81 |
| 100°C | 0.94 | 0.78 | 0.76 | 1.16 | 1.12 | 1.13 | 1.29 | 1.22 | 1.19 | 1.80 | 1.71 | |

Glycerol 2-phosphate, Pd/C

| | 2 h | | | 4 h | | | 18 h | | | 72 h | | |
|---|---|---|---|---|---|---|---|---|---|---|---|---|
| 50°C | n.d. | n.d. | n.d. | 0.77 | 0.83 | | 0.88 | 0.90 | 0.92 | 1.62 | 1.82 | 1.74 |
| 80°C | n.d. | n.d. | n.d. | 0.55 | 0.60 | | 0.64 | 0.66 | | 1.04 | 1.06 | 0.95 |
| 100°C | n.d. | n.d. | n.d. | 0.41 | 0.43 | | 0.54 | 0.56 | 0.59 | 0.94 | 1.06 | |

Acetyl phosphate

| | 18 h | | |
|---|---|---|---|
| 25°C | 7.56 | 7.50 | 8.05 |

Glucose 6-phosphate

| | 18 h | |
|---|---|---|
| 40°C | 0.63 | 0.82 |

**Table S2:** Raw ESI-LC–MS data and corresponding calculated product yields obtained from reactions containing Phi (200 mM) and ribose, cytidine, and creatine (100 mM each).

| Educts | Catalyst | Temp. [°C] | Time [h] | RT = 2.5 min | RT = 2.6 min | RT = 2.8min | Konz. µM |
|---|---|---|---|---|---|---|---|
| Ribose, Phi | Pd/C [1.5 mmol] | 50 | 72 | 486643 | 14860250 | 3678596 | 271.79 |
| Ribose, Phi | Pd/C [1.5 mmol] | 30 | 72 | 2081326 | 30010016 | 10443178 | 607.64 |

| Educts | Catalyst | Temp. [°C] | Time [h] | RT = 3.8 min | RT = 4.1 min | RT = 4.2min | RT = 5.8 min | RT = 6.3min | Konz. µM |
|---|---|---|---|---|---|---|---|---|---|
| Cytidine, Phi | Pd/C [1.5 mmol] | 50 | 72 | 9448673 | | | 8413277 | 11032208 | 191.93 |
| Cytidine, Phi | Pd/C [1.5 mmol] | 30 | 72 | 7132515 | 2193698 | 1194181 | 17797500 | 1938576 | 365.96 |

| Educts | Catalyst | Temp. [°C] | Time [h] | RT = 2.7 min | Konz. µM |
|---|---|---|---|---|---|
| Creatine, Phi | Pd/C [1.5 mmol] | 60 | 72 | 415 | 0.000208 |
| Creatine, Phi | Pd/C [1.5 mmol] | 60 | 18 | 5606792 | 2.803396 |

**Table S3:** Raw ESI-LC–MS data and calculated product yields obtained from reactions containing AMP (100 mM) and Phi (200 mM).

| Catalyst | Temp. [°C] | Time [h] | ADP 5'5'<br>RT =4.2 min<br>Konz. µM | ADP 3'5'<br>RT = 4.6 min<br>Konz. µM | ADP 2'5'<br>RT =5.3 min<br>Konz. µM |
|---|---|---|---|---|---|
| 5% Pd-$Ni_3Fe$ [0.015 mmol] | 50 | 18 | 6.09 | 5.15 | 3.38 |
| 5% Pd-$Ni_3Fe$ [0.015 mmol] | 100 | 18 | – | – | – |
| 5% Pd-$Ni_3Fe$ [0.015 mmol] | 50 | 96 | 5.12 | 5.3 | 3.25 |
| 5% Pd-$Ni_3Fe$ [0.015 mmol] | 100 | 96 | – | 1.49 | 2.35 |
